# ExBoX: a simple Boolean exclusion strategy to drive expression in neurons

**DOI:** 10.1101/2020.07.28.224691

**Authors:** Teresa Ubina, Tyler Vahedi-Hunter, Will Agnew-Svoboda, Wenny Wong, Akshay Gupta, Vijayalakshmi Santhakumar, Martin M. Riccomagno

## Abstract

The advent of modern single-cell biology has revealed the striking molecular diversity of cell populations once thought to be more homogeneous. This newly appreciated complexity has made intersectional genetic approaches essential to understanding and probing cellular heterogeneity at the functional level. Here we build on previous knowledge to develop a simple AAV-based approach to define specific subpopulations of cells by Boolean exclusion logic (AND NOT). This Expression by Boolean Exclusion (ExBoX) system encodes for a gene of interest which is turned On by a particular recombinase (Cre or FlpO) and turned Off by another. ExBoX allows for the specific transcription of a gene of interest in cells expressing only the activating recombinase, but not in cells expressing both. We show the ability of the ExBoX system to tightly regulate expression of fluorescent reporters *in vitro* and *in vivo*, and further demonstrate the adaptability of the system by achieving expression of a variety of virally-delivered coding sequences in the mouse brain. This simple strategy will expand the molecular toolkit available for cell- and time-specific gene expression in a variety of systems.

**Summary statement:** Ubina et al. describe the generation of a novel AAV-based intersectional approach to define and target specific subpopulations of cells in time and space via Expression by Boolean Exclusion (ExBoX).

## Introduction

Advancements in our understanding of the mechanisms underlying biological processes have been greatly dependent on the development of new genetic tools. A big breakthrough in mammalian genetics was the discovery and implementation of homologous recombination to generate loss-of-function alleles in mice (Capecchi, 1989). The introduction of recombinases as genetically encoded tools, in combination with conditionally targeted genetic alleles, made the control of genetic deletion in a tissue-specific manner possible (Gu et al, 1994; Tsien et al, 1996). With the availability of recombinase-dependent systems, region-specific gene knockouts and progenitor tracing have now become routine experimental strategies in mouse genetics (Branda & Dymecky, 2004).

Although cell diversity has always been appreciated in biology, single-cell profiling has revolutionized the way we think about certain organs by uncovering cell heterogeneity in populations that were once thought to display less complexity (Darmanis et al, 2015). Our ability to appreciate cellular complexity is currently limited by the available tools to label and manipulate cells with increased specificity. One way to overcome this limitation is to simply expand the pool of tissue- and population-specific recombinase lines. This requires the generation and characterization of independent transgenic recombinase mouse lines for each newly discovered cell population. A complementary approach that can take advantage of preexisting transgenic lines is to incorporate genetically encoded intersectional approaches (Awatramani et al, 2003). In comparison to conventional transgenic strategies that select cellular targets based on expression of one particular promoter or driver, intersectional approaches provide tighter specificity by selecting targets based on the overlapping or sequential expression of multiple recombinases (Andersson-Rolf et. al., 2017; Jensen & Dymecki, 2014; Robles-Oteiza et. al., 2015). These intersectional strategies based on the combinatorial expression of two recombinase systems (Cre/lox and Flp/FRT) were first used in the mouse to perform fate mapping of previously elusive neural progenitors (Dymecki, Ray, & Kim 2010). Intersectional approaches are not only able to label subpopulations with greater specificity, but also make a variety of Boolean logic operations (AND, NOT, OR) available (Daigle et al., 2019; Plummer et al, 2015). With multiple recombinase systems driving the expression of reporter genes, targeting of subpopulations of cells can be achieved through intersectional (expressing all drivers) and subtractive (lacking expression of one or more drivers) strategies (Farago et al, 2006).

Viral vectors offer an alternative approach to driving conditional and intersectional expression (Schnütgen et al, 2003; Atasoy et al, 2008; Gradinaru, 2010; Fenno et al, 2014). Strictly genetically encoded strategies have clear advantages over surgically introduced viral vectors, like non-invasiveness and broad applicability in hard-to-reach tissues during otherwise inaccessible embryonic stages (Jensen & Dymecki, 2014). However, viral expression systems can be extremely powerful given the ease of generation and the additional level of control one can gain by selecting the region and time of injection (Zhang et al, 2007; Sternson et al, 2015). There are now multiple versions of recombinase-dependent AAVs for recombinases like Cre, Flp, and DRE (Atasoy et al, 2008; Saunders et al, 2012; Xue, Atallah, & Scanziani, 2014). More recently, systems for intersectional expression driven by AAVs have also been developed (Kakava-Georgiadou et al, 2019; Fenno et al, 2014; Fan et al, 2020). Combination of these viral and genetically-encoded approaches will ultimately be essential to understand whether newly discovered molecular diversity within cell populations amounts to any notable phenotypic difference in terms of cellular structure and function.

In addition to being able to parse out functional diversity, there is an interest in the field of developmental biology in being able to specifically turn genes On and Off during defined developmental windows or critical periods (Wiesel & Hubel,1963; Kozorovitskiy et al, 2012). Intersectional approaches using Boolean negation/exclusion could provide a solution to this problem: a gene of interest could be turned On by a particular recombinase at a particular time point, while turned Off later (AND NOT) by a different recombinase (Table S1). Although a sophisticated system for AAV-driven expression using exclusion logic has been generated elsewhere (Fenno et al, 2014), the existing vectors have limitations in terms of being unnecessarily long and intricate in design, somewhat restricting their applicability. In this study we describe the design and characterization of an alternative expression system governed by Boolean exclusion logic and driven by a single AAV. The newly developed vectors are simpler in design and smaller in size, allowing for expression of longer genes of interest (GOI). As proof of principle, we validated the system in neuronal populations, and also generated tools for regulating neuronal activity in vivo.

## Results

### Design and construction of a combinatorial expression system using Boolean exclusion: CreOn-FlpOff ExBoX

To control expression and label subsets of cells with greater specificity using Cre and Flp recombinases, we set out to develop a simple AAV-based expression system governed by Boolean Exclusion logic (Expression by Boolean Exclusion or ExBoX). We first generated a system in which expression of a coding sequence of interest is turned On by Cre recombinase and turned Off by Flp recombinase: CreOn-FlpOff (Table S1). The CreOn-FlpOff vector was designed to contain a cassette with a coding sequence (CDS) of interest in-frame with a hemagglutinin tag (HA), P2A site, and EGFP reporter inverted with respect to a promoter. The promoter chosen for these experiments was that of human Synapsin I, which is a well characterized postmitotic neuronal promoter (Glover et al., 2002). The inverted cassette is flanked by Frt sites, and double-lox sites (lox2722 and loxP) in inverse orientation (DIO) (Schnütgen et al., 2003) (Figure 1A). Upon Cre-mediated recombination, an HA-tagged coding sequence of interest and EGFP get locked into the forward orientation, allowing for robust Cre-dependent expression in neurons. Conversely, in the presence of Flp the cassette is excised, resulting in abolishment of CDS and EGFP reporter expression (Figure 1A). Thus, the CDS is expressed only in the presence of Cre AND NOT Flp (Table S1).

**Figure 1.**
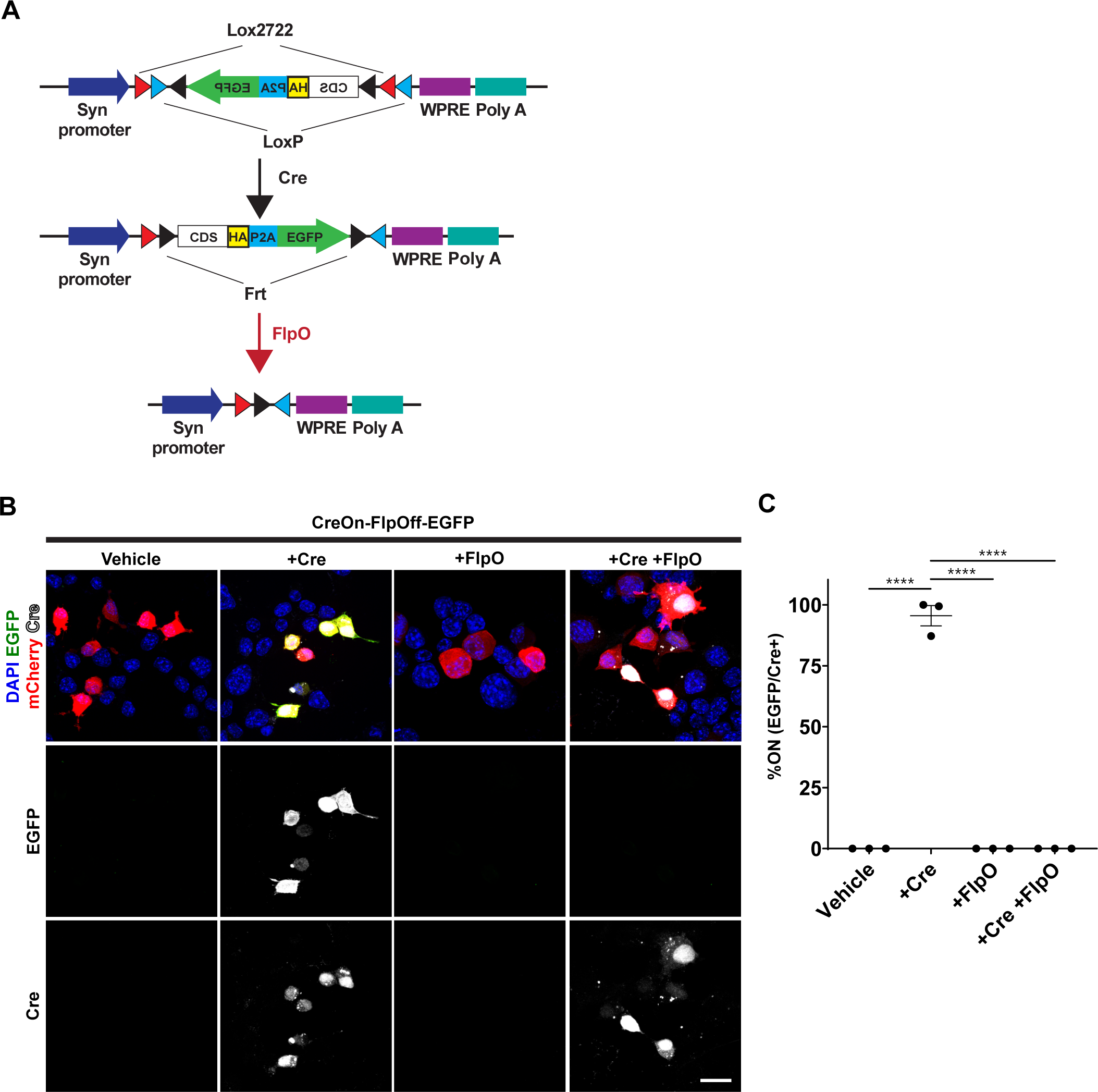
Design and proof of concept for CreOn-FlpOff constructs. **(A)** Schematic representation of the CreOn-FlpOff construct. It is composed of an inverted expression cassette encoding for a coding sequence (CDS) fused to an HA tag, a P2A site, and EGFP, which is flanked by Frt, lox2722, and loxP sites. Presence of Cre recombinase results in inversion of the cassette and expression. Presence of FlpO results in removal of the cassette. **(B)** Expression of fluorescent reporter proteins in Neuro2A cells transfected with CreOn-FlpOff-EGFP construct (green) and pCAG-mCherry (red, loading control). Co-transfection with pCAG-Cre plasmid causes inversion of the cassette into the “On” position and EGFP expression. Co-transfection of pCAG-FlpO deletes the cassette and abolishes EGFP expression. Fluorescent signals have been amplified in this image. Unamplified images can be seen in Figure S1A. **(C)** Quantification of Neuro2A cell transfections. Upon transfection with Cre, the CreOn-FlpOff-EGFP construct results in 95.5%±4.2 of Cre positive cells expressing EGFP, while co-transfection with FlpO results in abolishment of EGFP expression. Values given are mean±SEM. N=3 independent experiments, error bars ±SEM. ANOVA p<0.0001; Tukey post-hoc test: **** p<0.0001. Scale bar 30 μm Flp recombinase can be seen in Figure S1B.

For proof of concept, we first generated a plasmid carrying only EGFP in the reversible cassette of the CreOn-FlpOff system. The CreOn-FlpOff-EGFP plasmid was co-transfected with pCAG-IRES-Cre recombinase, pCAG-mCherry-IRES-FlpO recombinase, or both pCAG-IRES-Cre and pCAG-mCherry-IRES-FlpO in a neuroblastoma-derived cell line, Neuro2A (Figure 1B). mCherry-expressing plasmid was used as a transfection control when pCAG-mCherry-IRES-FlpO was not transfected. Upon transfection of the CreOn-FlpOff-EGFP with pCAG-mCherry control plasmid alone, no EGFP expression is found in any cells. As expected, when CreOn-FlpOff-EGFP plasmid is co-transfected with pCAG-IRES-Cre, the EGFP cassette is inverted to the correct orientation and 95.5%±4.2 of Cre positive cells express EGFP (Figure 1C). When CreOn-FlpOff-EGFP plasmid was co-transfected with pCAG-mCherry-IRES-FlpO, or co-transfected with pCAG-IRES-Cre and pCAG-mCherry-IRES-FlpO, no cells expressed EGFP, suggesting that FlpO can efficiently turn off reporter expression (Figure 1B,C). Similar results were observed by direct fluorescence when quantifying the EGFP^+^/mCherry^+^ cell ratio, or by staining for FlpO recombinase (Figure S1). This confirms that expression from CreOn-FlpOff ExBoX vectors can be successfully manipulated through Boolean exclusion in Neuro2A cells.

### Design of a complementary system that is turned On by Flp and turned Off by Cre: FlpOn-CreOff ExBoX

We next generated a complementary system that is turned On by Flp and Off by Cre, FlpOn-CreOff (Table S1). FlpOn-CreOff was designed using a similar internal logic, where an expression cassette containing a CDS, an in-frame HA tag, P2A site, and TdTomato reporter, are inverted with respect to the promoter. This expression cassette is flanked by lox2722 sites, and a Flp controlled DIO switch (fDIO or fFLEX) (Xue et al, 2014) containing F14 and Frt sites (Figure 2A). In this system, Flp inverts the cassette, resulting in CDS and TdTomato reporter expression, while Cre excises the cassette, causing the abolishment of CDS and TdTomato reporter expression (Figure 2A). Thus, CDS expression should only occur when FlpO is present AND NOT Cre (Table S1).

**Figure 2.**
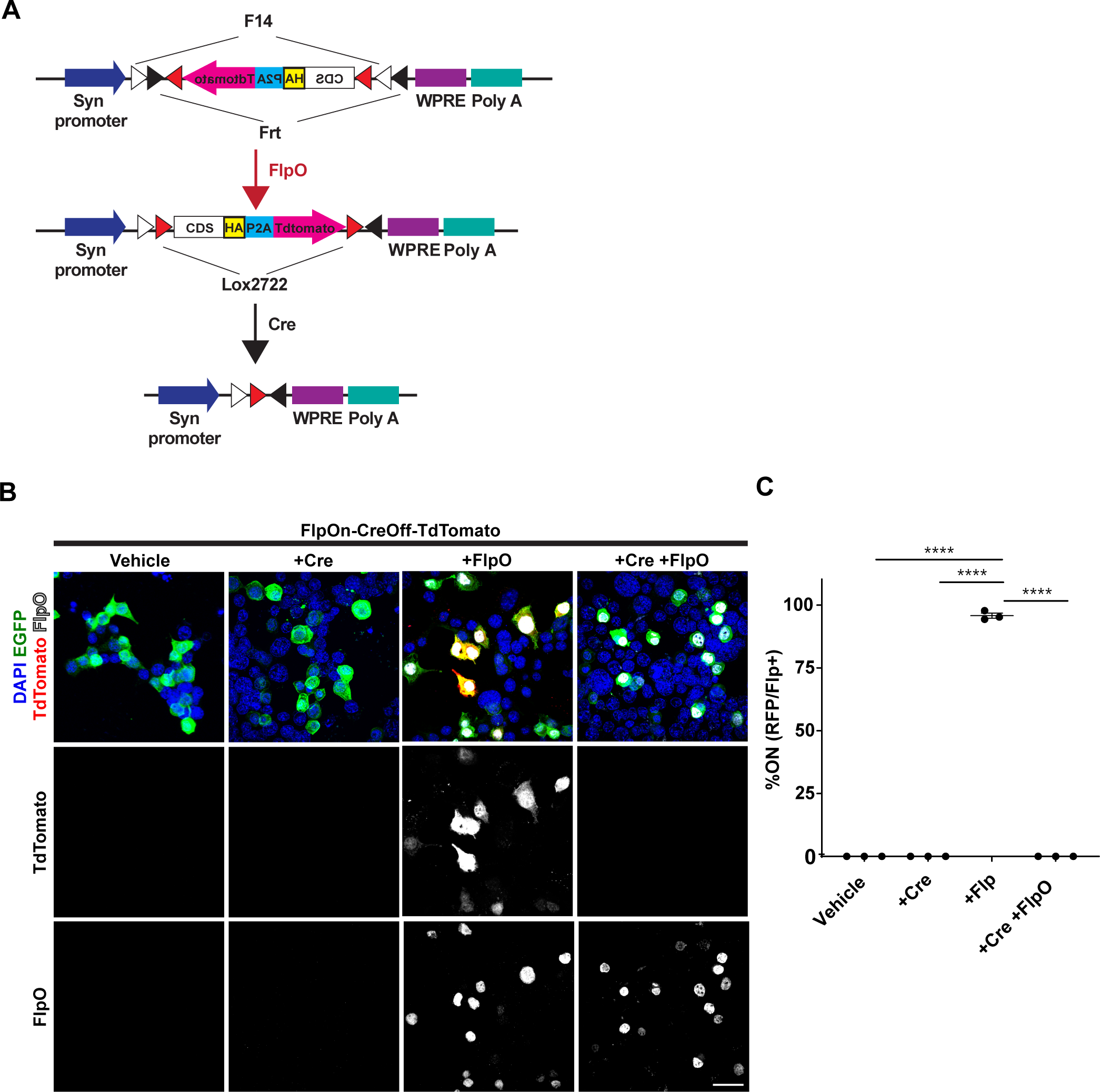
Design and proof of principle for FlpOn-CreOff constructs. **(A)** Schematic representation of the FlpOn-CreOff construct. The inverted expression cassette is flanked by lox2722, F14, and Frt sites. Presence of Flp recombinase results in inversion of the cassette and expression. Presence of Cre results in removal of the cassette. **(B)** Expression of fluorescent reporters in Neuro2A cells transfected with FlpOn-CreOff-TdTomato construct (red) and pCAG-EGFP (green, transfection control) in Neuro2A cells. Co-transfection with pCAG-FlpO causes inversion of the cassette into the “On” position and TdTomato expression. Transfection with pCAG-Cre removes the cassette and abolishes TdTomato expression. Fluorescent signals have been amplified in this image. Unamplified images can be seen in Figure S2A. **(C)** Upon transfection with FlpO, the FlpOn-CreOff-TdTomato construct results in detectable expression of reporter in 95.8%±1.0 of Flp positive cells, while co-transfection with Cre results in abolishment of TdTomato expression. Values given are mean±SEM. N=3 independent experiments, error bars ±SEM. ANOVA p<0.0001; Tukey post-hoc test: **** p<0.0001. Scale bar 30 μm. Antibody staining for Cre recombinase can be seen in Figure S2B.

As an initial proof-of-principle design, a FlpOn-CreOff vector driving simply a TdTomato fluorescent reporter was generated. FlpOn-CreOff-TdTomato plasmid was co-transfected with pCAG-Cre-IRES-EGFP, pCAG-FlpO, or both pCAG-Cre-IRES-EGFP and pCAG-FlpO. In the absence of pCAG-Cre-IRES-EGFP, a pCAG-IRES-EGFP plasmid was used as transfection control. When the FlpOn-CreOff-TdTomato plasmid was co-transfected with pCAG-FlpO, approximately 95.8%±1.0 of Flp positive cells displayed TdTomato expression (Figure 2B,C). When FlpOn-CreOff-TdTomato plasmid was co-transfected with pCAG-Cre-IRES-EGFP alone, there were no TdTomato expressing cells. Finally, as expected, co-transfection of the FlpOn-CreOff-TdTomato plasmid with both pCAG-Cre-IRES-EGFP and pCAG-FlpO resulted in no TdTomato reporter expression, confirming that Cre recombinase can abolish expression of this construct even in the presence of Flp (Figure 2B,C). Similar results were seen with direct fluorescence, and by Cre recombinase immunocytochemistry (Figure S2). Therefore, expression of the construct occurs only in the presence of Flp AND NOT Cre (Table S1).

### Characterization of the ExBoX system in neurons

To test whether this newly developed system was able to drive expression in neurons, we tested these constructs in primary cortical cultures. We used the same plasmid combinations that were used in Neuro2A cells to perform ex-utero electroporation prior to plating. Primary neuronal cultures were observed at 3 days in vitro (DIV). Transfection of CreOn-FlpOff-EGFP plasmid with pCAG-mCherry resulted in no reporter expression (Figure 3A). Notably, co-transfection of CreOn-FlpOff-EGFP plasmid with pCAG-IRES-Cre resulted in 100%±0.0 of transfected neurons expressing EGFP as expected (Figure 3B). When CreOn-FlpOff-EGFP plasmid was co-transfected with pCAG-mCherry-IRES-FlpO or with pCAG-mCherry-IRES-FlpO and pCAG-IRES-Cre, we found no EGFP reporter expression in any transfected cells (Figure 3A,B).

**Figure 3.**
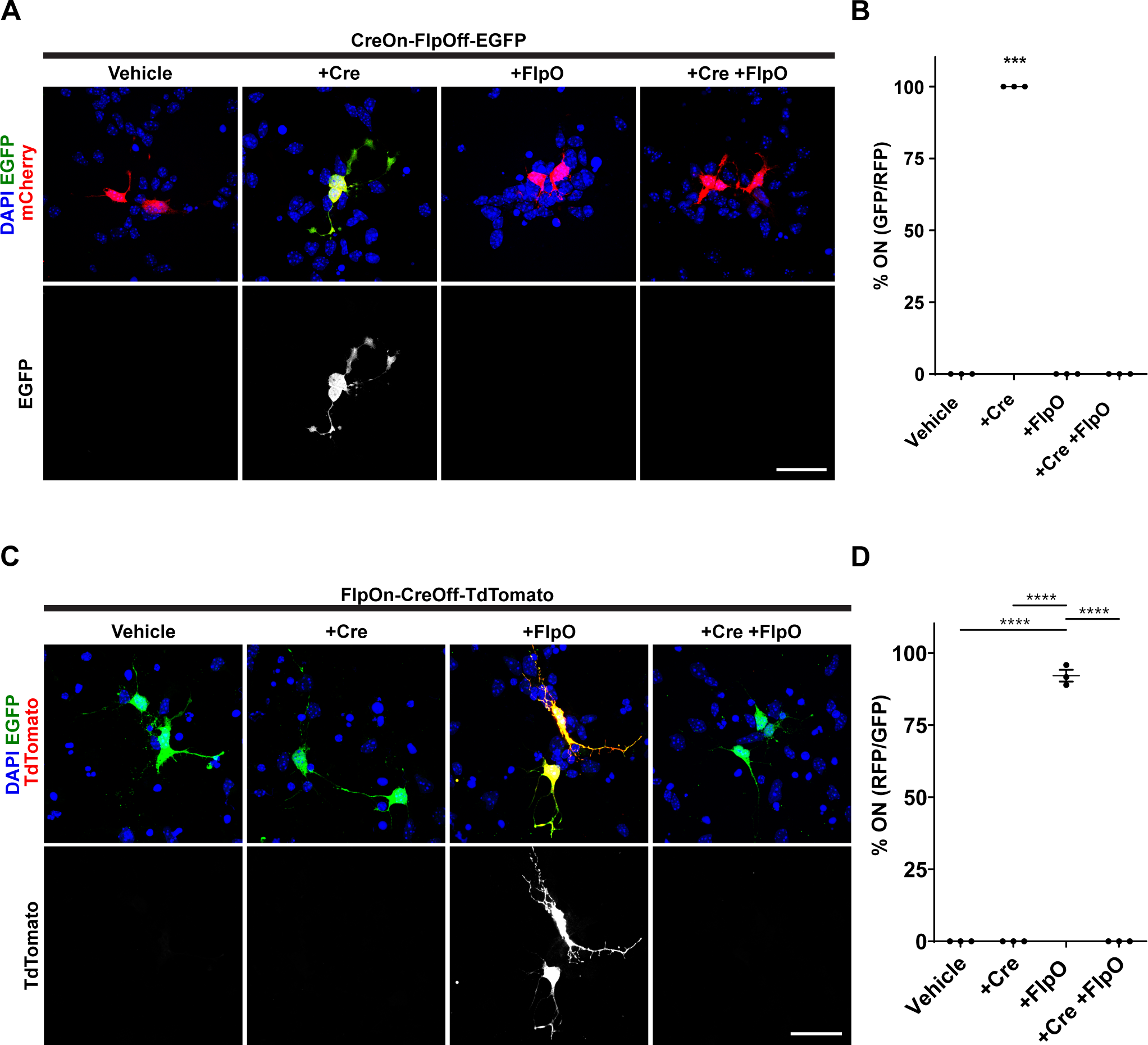
Validation of ExBoX in primary cortical neurons. **(A)** Primary cortical neurons transfected with CreOn-FlpOff-EGFP and either pCAG-IRES-Cre, pCAG-mCherry-IRES-FlpO, of both pCAG-IRES-Cre and pCAG-mCherry-IRES-FlpO. **(B)** Cortical neurons transfected with CreON-FlpOff-EGFP and pCAG-IRES-Cre results in 100%±0.0 of transfected cells expressing EGFP. Without Cre of in the presence of FlpO there is no EGFP expression in any neurons. pCAG-mCherry was co-transfected as transfection control. Values given are mean±SEM. N=3 independent experiments, error bars ±SEM. Fisher’s exact test ***p<0.0001. **(C)** Primary cortical neurons transfected with FlpOn-CreOff-TdTomato and pCAG-FlpO, pCAG-Cre-IRES-EGFP, or both pCAG-FlpO and pCAG-Cre-IRES-EGFP. **(D)** Cortical neurons transfected with FlpOn-CreOff-TdTomato and FlpO results in detectable TdTomato expression in 92.1%±2.0 of transfected cells. pCAG-EGFP was co-transfected as transfection control. Without FlpO, or in the presence of Cre, there is no TdTomato expression in any neurons. ANOVA p<0.0001; Tukey post-hoc test: **** p<0.0001. Fluorescent signals have been amplified in all images. Scale bar 30 μm

Similarly, transfection of FlpOn-CreOff-TdTomato plasmid with EGFP vehicle control resulted in no reporter expression (Figure 3C). Co-transfection of FlpOn-CreOff-TdTomato plasmid with pCAG-FlpO and pCAG-IRES-EGFP resulted in 92.1%±2.0 of transfected cells expressing the TdTomato reporter, as expected (Figure 3D). If FlpOn-CreOff-TdTomato plasmid is co-transfected with pCAG-Cre-IRES-EGFP alone, or pCAG-Cre-IRES-EGFP and pCAG-FlpO, it is expected that there will be no TdTomato expression since Cre should excise the cassette. As predicted, no cells transfected in this combination had any TdTomato expression (Figure 3C,D).

We then tested the neuronal expression of the ExBoX system *in vivo.* Initial validation of CreOn-FlpOff was performed by *in utero* electroporation (IUE) of CreOn-FlpOff-EGFP plasmid with pCAG-mCherry vehicle, pCAG-IRES-Cre and pCAG-mCherry, pCAG-mCherry-IRES-FlpO alone, or with pCAG-IRES-Cre and pCAG-mCherry-IRES-FlpO into the lateral ventricle of E15.5 mice, followed by visualization 48 hours later. When the CreOn-FlpOff-EGFP plasmid was electroporated with mCherry vehicle alone or with pCAG-mCherry-IRES-FlpO, we found no EGFP expression (Figure 4A). Tissue co-electroporated with CreOn-FlpOff-EGFP plasmid, pCAG-IRES-Cre, and pCAG-mCherry had robust expression of the EGFP reporter, resulting in labeling of 97.7%±0.9 of transfected cells (Figure 4A,B). Co-electroporation of CreOn-FlpOff-EGFP plasmid with pCAG-IRES-Cre and pCAG-mCherry-IRES-FlpO resulted in negligible EGFP expression (1.7%±0.9 of transfected cells, Figure 4A,B).

**Figure 4.**
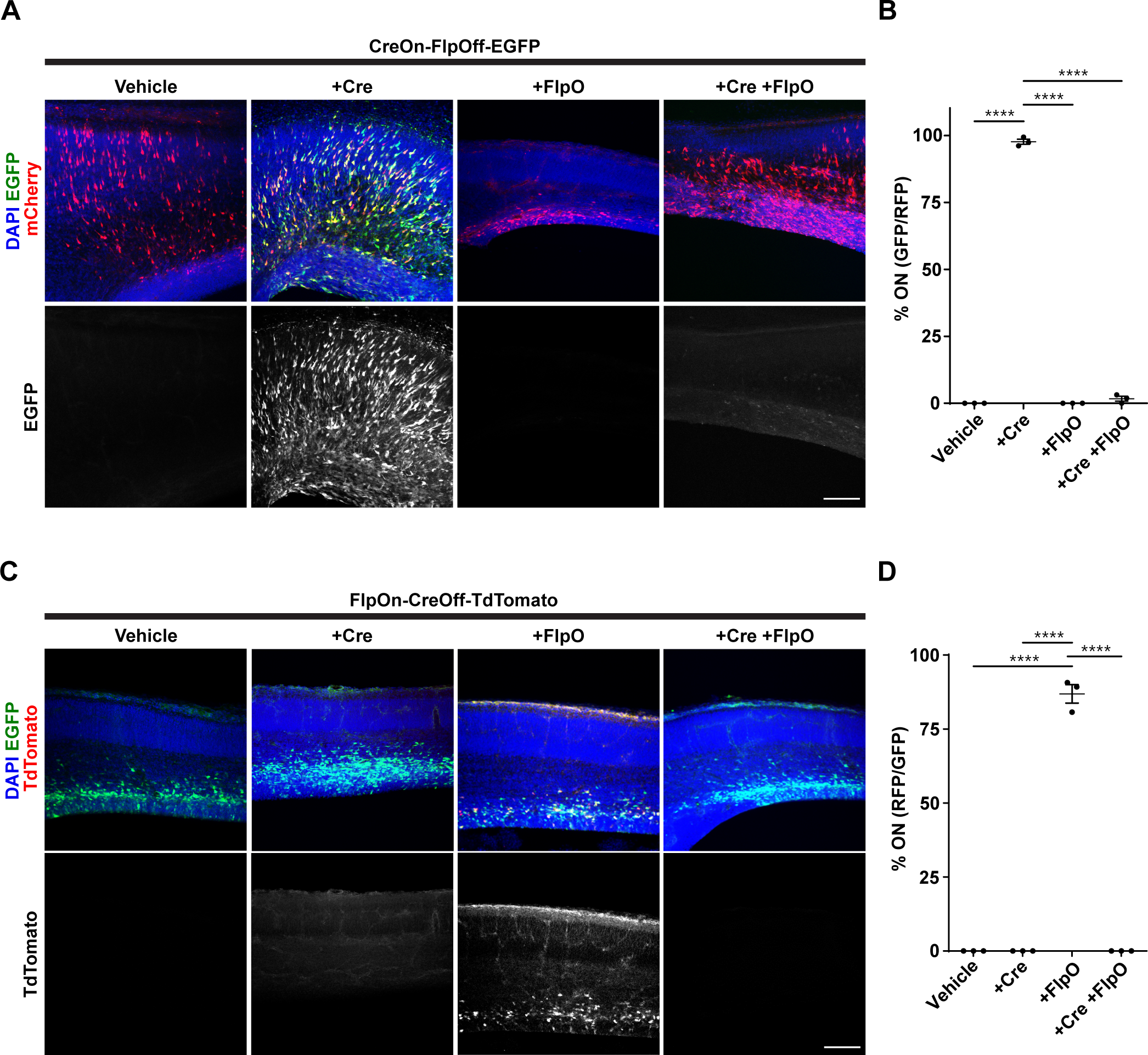
*In vivo* validation of CreOn-FlpOff and FlpOn-CreOff vectors via *in utero* electroporation (IUE). **(A)** Co-electroporation of CreOn-FlpOff-EGFP with mCherry reporter, pCAG-IRES-Cre, pCAG-mCherry-IRES-FlpO, or pCAG-IRES-Cre and pCAG-mCherry-IRES-FlpO in E15.5 ICR mouse embryos. **(B)** Electroporation of CreOn-FlpOff-EGFP with pCAG-IRES-Cre results in 97.7%±0.9 of transfected cells expressing EGFP. Electroporation with pCAG-mCherry-IRES-FlpO results in no expression and electroporation with pCAG-IRES-Cre and pCAG-mCherry-IRES-FlpO results in only 1.7%±0.9 of cells still expressing EGFP. mCherry expression serves as electroporation control. **(C)** Co-electroporation of FlpOn-CreOff-TdTomato with pCAG-IRES-EGFP, pCAG-Cre-IRES-EGFP, pCAG-FlpO, or pCAG-Cre-IRES-EGFP and pCAG-FlpO in E15.5 ICR embryos. pCAG-IRES-EGFP expression was used as electroporation control. **(D)** Electroporation of FlpOn-CreOff-TdTomato with pCAG-FlpO results in 86.9%±3.1 of transfected cells expressing TdTomato. Co-electroporation with pCAG-Cre-IRES-EGFP, or with pCAG-Cre-IRES-EGFP and pCAG-FlpO, results in no cells expressing TdTomato. Values given are mean±SEM. N=3 individual animals, error bars ±SEM. ANOVA p<0.0001 for both sets of experiments; Tukey post-hoc test: **** p<0.0001 vs. all other transfections. Fluorescent signals have been amplified. Scale bar 100 μm.

To validate FlpOn-CreOff, we electroporated the FlpOn-CreOff-TdTomato plasmid alone and this resulted in no TdTomato expression. A plasmid driving EGFP under the pCAG promoter was used as electroporation control. Co-electroporation of FlpOn-CreOff-TdTomato plasmid with pCAG-FlpO resulted in inversion of the cassette and detectable expression of TdTomato in 86.9%±3.1 of transfected cells (Figure 4C,D). Co-electroporation of FlpOn-CreOff-TdTomato plasmid with pCAG-Cre-IRES-EGFP showed no TdTomato expression (Figure 4C,D). Finally, no TdTomato expression was detected when the FlpOn-CreOff-TdTomato plasmid was co-electroporated with pCAG-FlpO and pCAG-Cre-IRES-EGFP, indicating successful AND NOT exclusion (Figure 4C,D). These data suggest that FlpOn-CreOff and CreOn-FlpOff ExBoX work efficiently in neurons, *in vitro* and *in vivo*.

### Validation of ExBoX AAVs *in vivo*

To generate tools that would be useful for manipulation of gene expression in the postnatal brain, we packaged the FlpOn-CreOff construct into AAV9. To validate that expression of the FlpOn-CreOff constructs is maintained in the virally-packaged form, we performed stereotactic injections to deliver FlpOn-CreOff-TdTomato AAV into the dentate gyrus (DG) of adult mice *in vivo*. FlpOn-CreOff-TdTomato AAV was co-injected with either AAV-EF1α-FlpO-WPRE (AAV-EF1α-FlpO), or with AAV-EF1α-FlpO and pENN.AAV.CamKII 0.4. Cre. SV40 (AAV-CamkII-Cre), and expression was observed after 3 weeks (Figure 5A-I). AAV expressing EGFP was co-injected in wild type animals and when Cre was injected to visualize the injection site. Similar to our previous findings with FlpOn-CreOff plasmids, robust TdTomato reporter expression was observed in DG neurons co-infected with AAV-FlpOn-CreOff-TdTomato and AAV-EF1α-FlpO (Figure 5B,E), verifying that expression of the construct in the presence of the ‘On’ recombinase was maintained in viral form. Furthermore, AAV-FlpOn-CreOff-TdTomato reporter expression was inactivated by AAV-CamkII-Cre even in the presence of AAV-EF1α-FlpO, with only 1.5%±0.9 of DG cells co-expressing TdTomato and EGFP (Figure 5C, F, I). Thus, the FlpOn-CreOff construct works as expected when packaged into AAV9.

**Figure 5.**
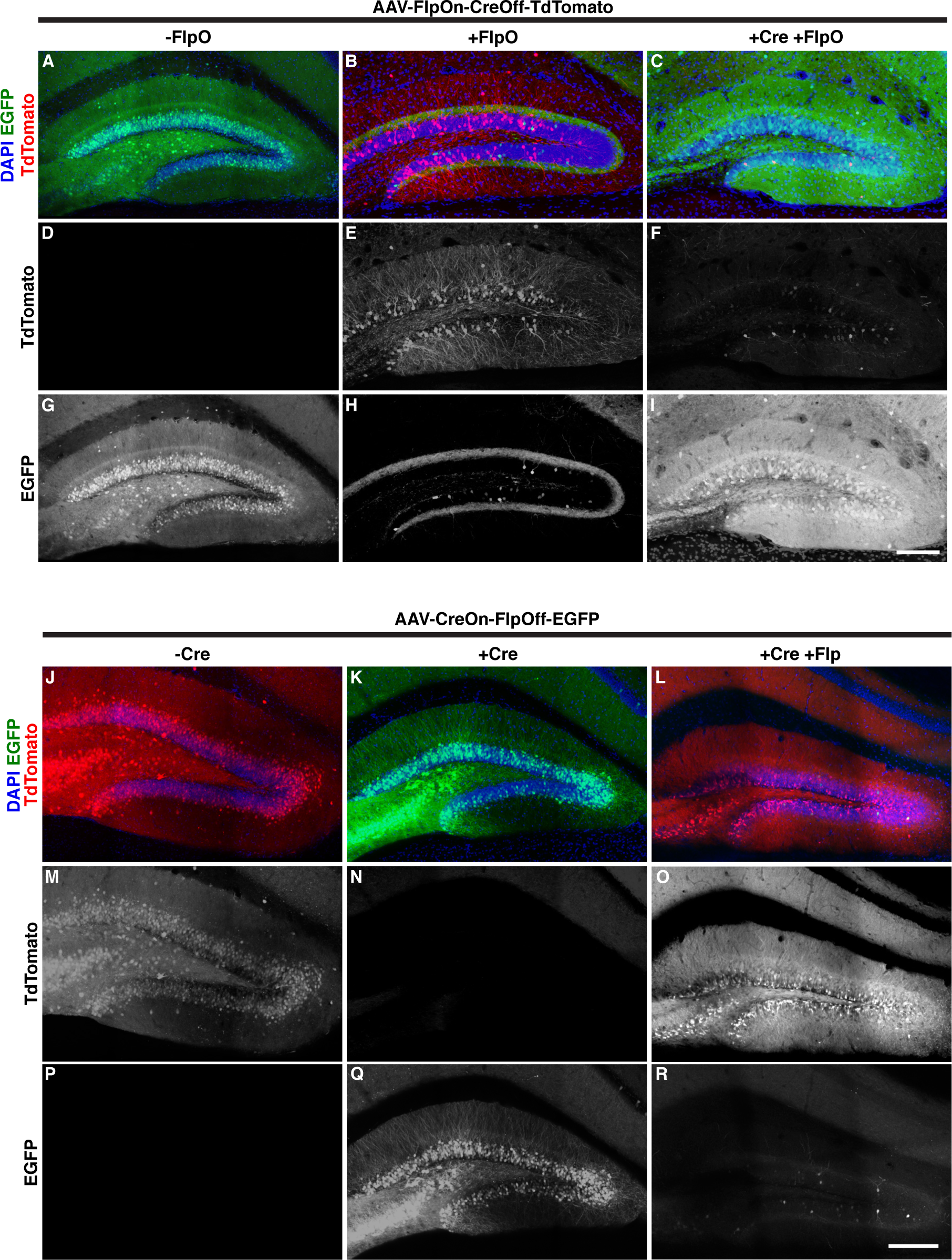
*In-vivo* validation of FlpOn-CreOff and CreOn-FlpOff AAV by stereotactic injection in postnatal DG neurons. **(A-I)** Expression of viral FlpOn-CreOff-TdTomato construct (red) in WT animals co-injected with pAAV-CAG-GFP **(A, D, G)**, with AAV-EF1α -FlpO-WPRE **(B, E, H),** or AAV-EF1α -FlpO-WPRE + AAV-CamKII-Cre + AAV-CAG-FLEX-EGFP **(C, F, I)**. EGFP expression is used to visualize the injection site in the absence of TdTomato expression. Even in the presence of FlpO, Cre injection resulted in inactivation of TdTomato reporter expression (**C**, **F**). DAPI (blue) was used to visualize nuclei. Expression of viral CreOn-FlpOff-EGFP construct (green) in WT animals (**J, M, P**), in Grik4-Cre animals (**K, N, Q**), and in Grik4-Cre mice co-injected with AAV-EF1α -FlpO-WPRE (**L, O, R**). pAAV-CAG-TdTomato (red) was co-injected when necessary to visualize the injection site. Even in the presence of Cre, FlpO injection resulted in inactivation of reporter expression by the CreOn-FlpOff-EGFP virus (**L, O, R**). EGFP labeling in the molecular layer of AAV-EF1α -FlpO injected animals (**L, O**) is from axonal projections from the contralateral side. DAPI (blue) was used to visualize nuclei. Image fluorescence is unamplified except for EGFP in J-R. Scale Bar 150 μm

We next packaged the complementary CreOn-FlpOff construct into AAV9. To validate that expression of the CreOn-FlpOff construct is maintained in the virally-packaged form, we performed stereotactic injections to deliver CreOn-FlpOff-EGFP AAV into the dentate gyrus (DG) of adult mice *in vivo*. For validation for the viral CreOn-FlpOff construct we incorporated a genetically-encoded Cre recombinase driven by a transgene. The *Grik4-cre* mouse line was specifically chosen for providing robust Cre expression in the hippocampus, including the DG (Figure S3). AAV-CreOn-FlpOff-EGFP was injected independently into *Grik4-Cre* mice or Cre-negative littermates (WT), or co-injected with AAV-EF1α-FlpO into *Grik4-Cre* animals (Figure 5J-R). AAV9 constitutively expressing TdTomato (pAAV-CAG-TdTomato) was co-injected in WT animals and with AAV-EF1α-FlpO to visualize the injection site. When AAV-CreOn-FlpOff-EGFP was injected into WT (Cre-negative) animals, no EGFP reporter expression was observed (Figure 5J, P). Conversely, injections into *Grik4-Cre* positive animals demonstrated robust EGFP reporter expression (Figure 5K,Q), verifying the Cre-dependent activation of the virally-packaged construct. Furthermore, in *Grik4-Cre* animals co-injection of AAV-EF1α-FlpO resulted in efficient inactivation of EGFP reporter expression (2.8%±1.2 EGFP^+^;mCherry^+^ escaper cells; Figure 5L,O, R, S4). These results suggest that the virally-packaged ExBoX constructs are successful at driving gene expression through the expected AND NOT Boolean operation.

### Generation of Activity-modulating CreOn-FlpOff Viruses for Neuroscience

To further validate the ExBoX system using different cargos, we next generated CreOn-FlpOff AAV constructs co-expressing EGFP and well-characterized modulators of neuronal firing or synaptic release. To reduce activity, we generated a CreOn-FlpOff virus that expresses Kir2.1 (AAV-CreOn-FlpOff-Kir2.1-EGFP), an inward-rectifying K+ channel that has been demonstrated to increase rheobase current and reduce resting membrane potential of neurons when overexpressed (Xue et al, 2014). In parallel, we cloned a version of Kir2.1 carrying three point mutations that render the channel in a non-functional state, referred to as Kir2.1Mut (AAV-CreOn-FlpOff-Kir2.1Mut-EGFP), which can be used as a negative control for the Kir2.1 construct (Xue et al, 2014). We also designed a CreOn-FlpOff virus encoding mNaChBac (AAV-CreOn-FlpOff-mNaChBac-EGFP). This bacterial Na^+^ ion channel has been demonstrated to increase firing of neurons when overexpressed by reducing action potential threshold and rheobase (Xue et al, 2014). To serve as negative control for mNaChBac in neural circuit experiments, we cloned a non-conducting mNaChBac mutant referred to as mNaChBacMut into the CreOn-FlpOff vector (AAV-CreOn-FlpOff-mNaChBacMut-EGFP) (Xue et al, 2014). Finally, we generated CreOn-FlpOff encoding Tetanus Neurotoxin (AAV-CreOn-FlpOff-TeNT-EGFP) to block synaptic transmission (Sweeney et al, 1995).

We validated reporter expression of these five viral constructs in adult mice using stereotactic injection to deliver each independently or in combination with AAV-EF1 -αFlpO into the DG of adult Grik4-Cre mice or WT (Cre-negative) littermates (Figure 6). To visualize the injection site pAAV-CAG-TdTomato was co-injected in WT, or when AAV-EF1α-FlpO was injected. As expected, injection of any of the viral constructs into Cre-negative animals produced no EGFP reporter expression (Figure 6A,E,I,M,Q). Conversely, each virus was able to produce robust EGFP reporter expression in DG neurons when injected into Grik4-Cre positive animals (Figure 6B,F,J,N,R), demonstrating expression of these viral constructs is Cre-dependent. Furthermore, in *Grik4-Cre* mice, injection of AAV-EF1α-FlpO prevented EGFP reporter expression, with the exception of very few escaper cells (Figure 6C,D,G,H,K,L,O,P,S,T; Figure S4). Thus, despite the larger cargo size, expression of these activity-modulating CreOn-FlpOff viral constructs was similar to that of the CreOn-FlpOff-EGFP viral construct (Figure 5J-R, 6, S4).

**Figure 6.**
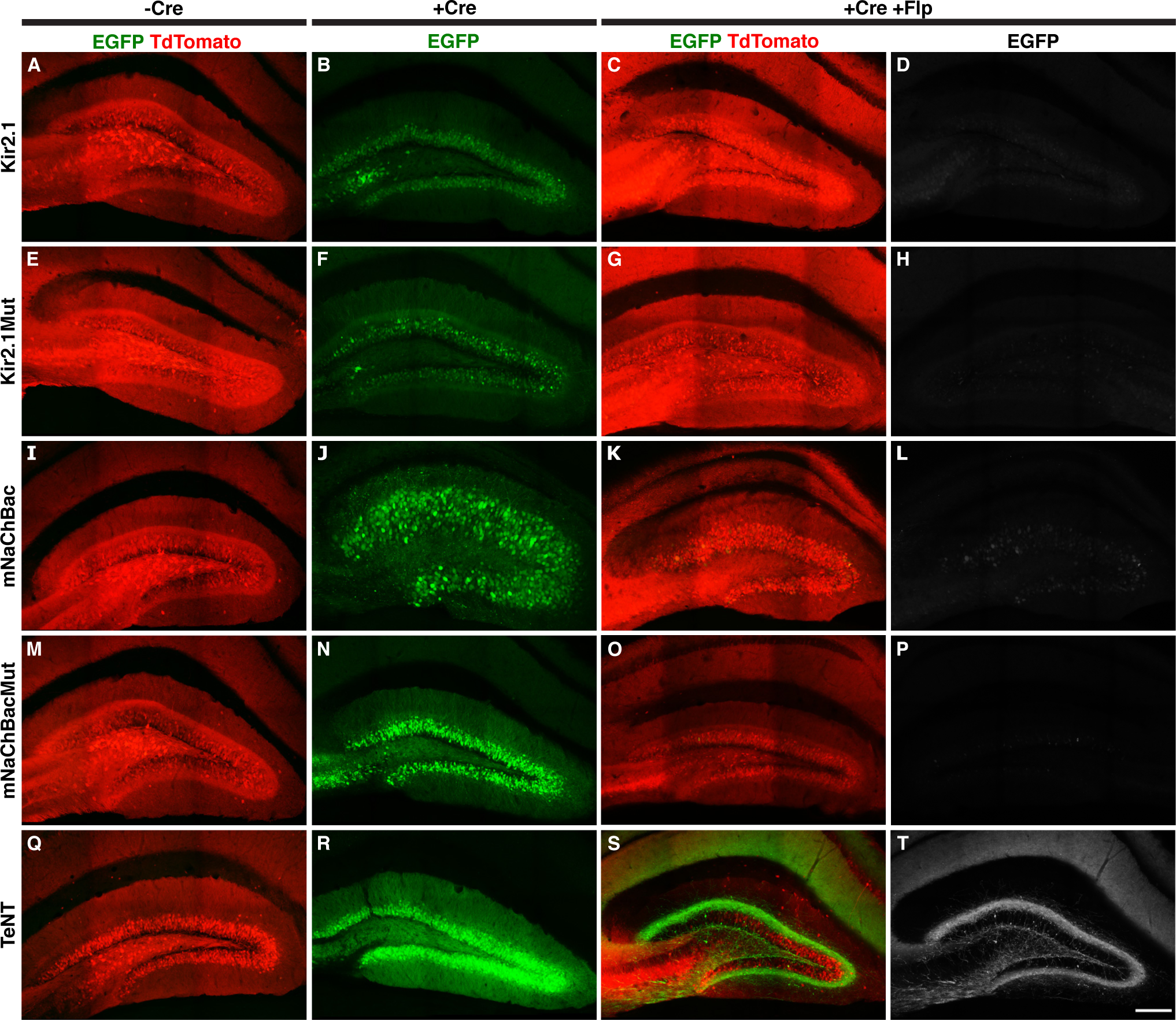
*In-vivo* validation of CreOn-FlpOff AAV constructs to manipulate neuronal activity. **(A-T)** Expression of fluorescent reporter by CreOn-FlpOff AAVs (green) co-expressing (A-D) Kir2.1, (E-H) Kir2.1-Mutant, (I-L) mNaChBac, (M-P) mNaChBac-mutant, and **(Q-T)** TeNT in WT (A, E, I, M, Q), *Grik4-Cre* animals **(B, F, J, N, R)** and *Grik4-Cre* animals co-injected with AAV-EF1α -FlpO-WPRE (C, D, G, H, K, L,O, P, S, T). Injections delivering AAV-EF1α -FlpO included pAAV-CAG-TdTomato (red) to visualize infected cells. In the presence of Cre and FlpO, expression of the CreOn-FlpOff virus was inactivated (C, D, G, H, K, L, O, P, S, T). EGFP labeling in the molecular layer of TeNT/FlpO injected animals (T) is from axonal projections from the contralateral side. EGFP fluorescence is amplified in all images. Scale bar 150 μm

### Temporal and spatial control of expression using ExBoX

So far our data suggest that the ExBoX system can be used to efficiently control expression of multiple genes of interest in a Cre- and Flp-dependent manner through AND NOT Boolean operations. Thus, this system has the potential to be a powerful tool to control expression temporally and spatially (Figure 7A,B). To test the usefulness of the ExBoX vectors for temporal control of expression, we established a simple *in vitro* paradigm. Neuro2A cells were transfected with FlpOn-CreOff-TdTomato, pCAG-FlpO, pCAG-ERT2-Cre-ERT2, and pCAG-EGFP. 48 hours after transfection cells were treated with 4-hydroxytamoxifen (4OHT) or vehicle three days in culture and collected for analysis four days later. Before treatment 87.7%±1.7 of transfected cells were TdTomato positive (On condition) (Figure 7C,D). In vehicle-treated cultures 96.1%±1.8 of transfected cells were positive for TdTomato, while only 3.2%±1.0 of 4OHT treated EGFP^+^ cells remained TdTomato positive, a highly significant decrease (Figure 7C,D). These results show robust temporal control of ExBoX-driven expression in Neuro2A cells.

**Figure 7.**
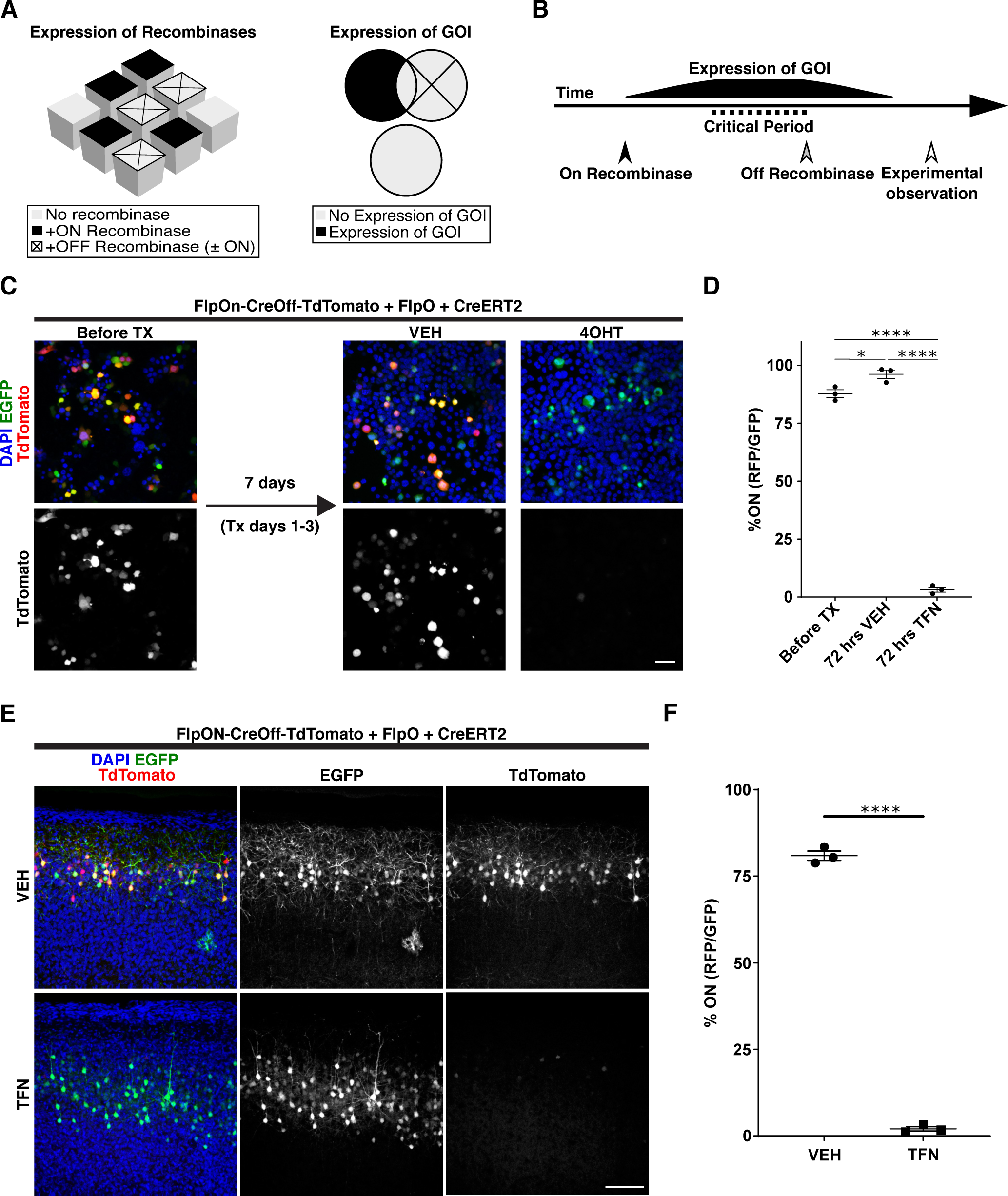
The ExBoX system can be utilized in multiple ways. **(A)** Expression of a gene of interest (GOI) can be restricted to a specific subpopulation of cells. Only the subpopulation of cells expressing the On recombinase, but not the Off recombinase, will express the GOI. **(B)** ExBoX can also be used for temporal control of expression. In this scheme, activation of the On recombinase is used just prior to a critical period of interest. Introduction or activation of the Off recombinase at the end of this critical period will turn off expression of the GOI. This serves to reduce confounding factors during later experimental observation, as GOI expression is no longer present. **(C)** Neuro2A cells were transfected with FlpOn-CreOff-TdTomato, FlpO, ERT2-Cre-ERT2, and EGFP. Beginning 48 hours after transfection, cultures were treated with 4-hydroxytamoxifen (4OHT) or vehicle (VEH) for three days and collected for imaging and quantification four days after that. **(D)** Before treatment 87.7%±1.7 of cells were On. After treatment with the vehicle 96.1%±1.8 of cells were On, while only 3.2%±1.0 of cells treated with 4OHT remained On. Values given are mean±SEM. N=3 independent experiments, ANOVA p<0.0001; Tukey post-hoc test: *p=0.0205, ****p<0.0001. Error bars ±SEM, scale bar 50μm. **(E)** Temporal control of FlpOn-CreOff-TdTomato expression *in* vivo. IUE was performed with the same combination of plasmids as in C. Pups were gavaged with oil or tamoxifen from P10-P13 and collected for visualization 10 days later. **(F)** In vehicle treated pups, 80.9%±1.4 of transfected cells co-expressed EGFP and TdTomato. Tamoxifen treatment resulted in a significant decrease of TdTomato expression in EGFP positive cells (2.1%±0.6; two-tailed unpaired t-test, N=3 independent injections, ****p<0.0001). Values given are mean±SEM. Image fluorescence is unamplified. Scale bar 100 μm

To further demonstrate the ability of ExBoX to provide temporal control of expression *in vivo,* we performed IUEs at E16.5 with FlpOn-CreOff-TdTomato in combination with pCAG-FlpO, pCAG-ERT2-Cre-ERT2, and pCAG-EGFP. After birth, pups were orally gavaged with tamoxifen or vehicle oil from P10 to P13 (4 days) and collected for visualization 10 days later (P23). Pups orally gavaged with vehicle showed 80.9%±1.4 of EGFP^+^ cells also expressing TdTomato (Figure 7E,F). Tamoxifen-treated pups displayed a significant decrease in EGFP^+^ cells expressing TdTomato, with only 2.1%±0.6 remaining in the On condition after treatment (Figure 7E,F). Taken together, these experiments corroborate that FlpOn-CreOff ExBoX constructs can be used in combination with tamoxifen-inducible Cre recombinase to control temporal expression of GOIs *in vitro* and *in vivo*.

Finally, we sought to demonstrate that ExBoX-driven expression can be temporally and spatially controlled by performing successive viral injections. ICR pups were initially injected into the lateral ventricles with AAV-CreOn-FlpOff-EGFP and AAV9-hSyn-Cre at Postnatal day 1 (P1) to obtain robust expression in Layer V of the cortex (Mathiesen, 2020). These animals were injected 3 weeks later (P22) with pAAV-CAG-TdTomato alone or in combination with AAV-EF1a-FlpO into the primary somatosensory cortex (S1). Tissue was collected for analysis 3 weeks after the P22 injection. Because the initial P1 injection robustly labels layer V, quantification was limited to this layer at the site of the second injection. Only injections that were surrounded by expression of the CreOn-FlpOff-EGFP construct on both sides were scored. We found that 46.8%±7.1 of labeled cells co-expressed EGFP and TdTomato when the control pAAV-CAG-TdTomato vector was injected (Figure 8A,B). Notably, co-injection of pAAV-CAG-TdTomato and pAAV-EF1α-FlpO resulted in a strong and significant decrease in expression, with only 7.6%±2.1 of the cells co-expressing EGFP and TdTomato (Figure 8A,B).

**Figure 8.**
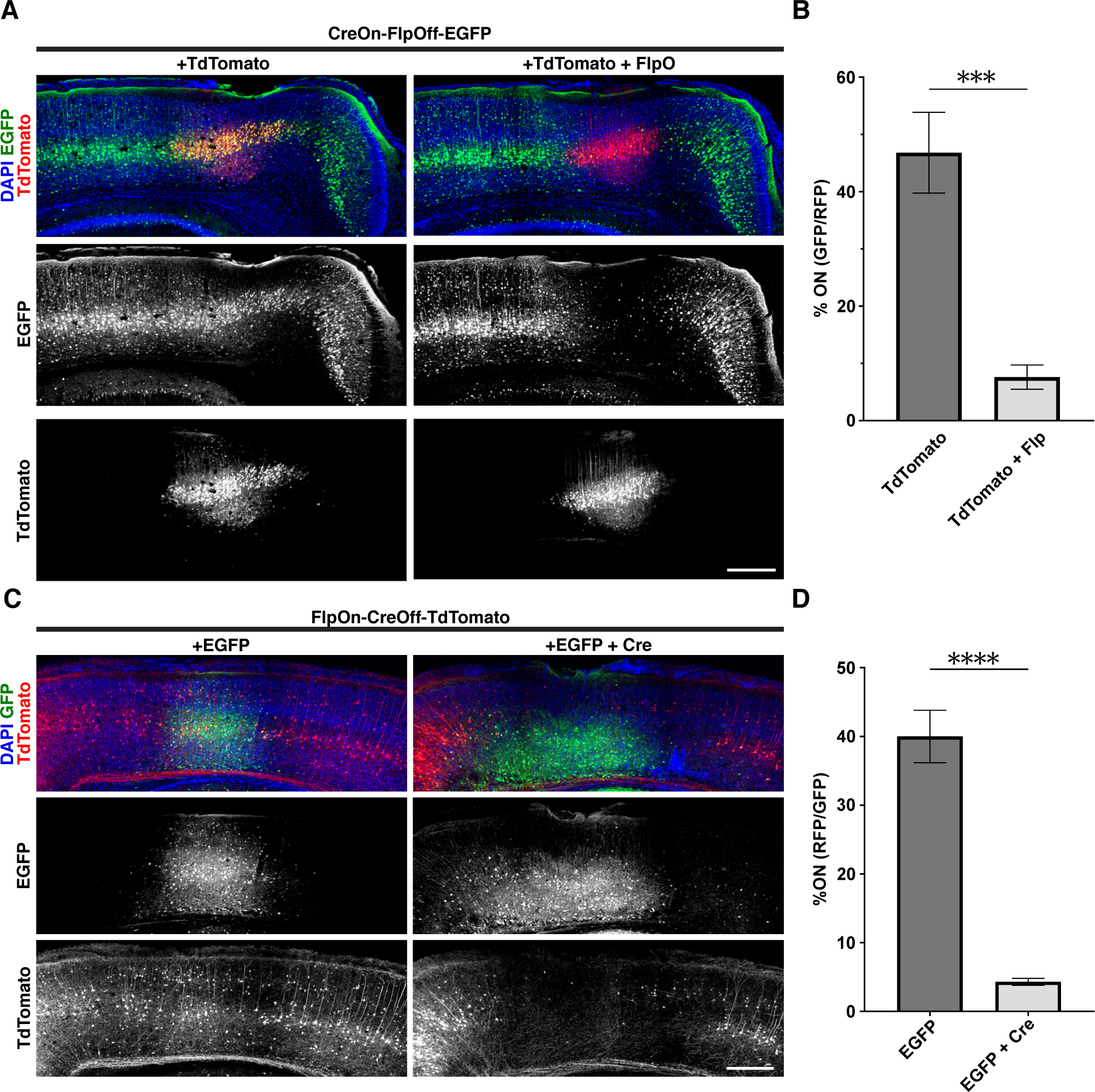
Spatial and temporal control of expression by ExBoX AAVs *in vivo*. **(A)** Detection of AAV-CreOn-FlpOff-EGFP co-injected intracerebroventricularly with AAV-hSyn-Cre at P1. pAAV-CAG-TdTomato alone or in combination with AAV-EF1α -FlpO into layer V of S1 at P22. (B) Quantification of A. 46.8%±7.1 of LV cells injected with pAAV-CAG-TdTomato alone express EGFP. A significant decrease in the number of EGFP-expressing cells is observed in animals co-injected with pAAV-CAG-TdTomato and AAV-EF1α -FlpO (7.6%±2.1; two-tailed unpaired t-test, N=5, ***p=0.0007). (C) AAV-FlpOn-CreOff-TdTomato expression when co-injected with AAV-EF1α -FlpO intracerebroventricularly at P1. pAAV-CAG-GFP alone or with AAV-hSyn-Cre into layer V of S1 at P22. (D) Injection of pAAV-CAG-GFP alone results in 40.0%±3.8 of cells at the injection site in the On condition (expressing TdTomato). A significant decrease of TdTomato-expressing cells (4.3%±0.5) at the injection site was observed when pAAV-CAG-GFP and AAV-hSYN-Cre were co-injected (two-tailed unpaired t-test, N=3 independent injections for EGFP control, N=5 independent injections for EGFP+Cre, ****p<0.0001). Error bars ±SEM. Scale bars 300 μm. Values given are mean±SEM.

A similar proof-of-principle experiment was performed to validate the temporal and regional control of expression of the FlpOn-CreOff AAV constructs. AAV-FlpOn-CreOff-TdTomato and AAV-EF1a-FlpO were co-injected into the lateral ventricles of P1 pups. Three weeks later, AAV-CAG-EGFP alone or in combination with AAV9-hSyn-Cre were injected stereotactically into S1. In animals injected with AAV-CAG-GFP alone, 40.0%±3.8 of Layer V cells co-expressed EGFP and TdTomato (Figure 8C,D). Animals injected with AAV-CAG-GFP and AAV9-hSyn-Cre showed a significant 9-fold decrease in the number of cells co-expressing EGFP and TdTomato (4.31%±0.5; Figure 8C,D). Overall, our data suggest that the ExBoX system can be used to provide temporal and spatial control of expression of genes of interest in a Cre- and Flp-dependent manner.

## Discussion

### Design of a simpler, more space-efficient Boolean exclusion expression system

The ability to identify and characterize specific cell types and subpopulations has been challenging in part because, until recently, cell targeting strategies have been limited to populations defined by expression of a single marker or reporter. In order to solve this issue and increase specificity of target selection, a number of intersectional strategies have been developed (Andersson-Rolf, 2017; Atasoy et al, 2008; Fenno et al, 2014; Gradinaru et al, 2010; Jensen & Dymecki, 2014; Plummer et al, 2015; Robles-Oteisa, 2015; Schnütgen et al, 2003). A previous study described the generation of an expression system using Boolean logical operations, all governed by a single AAV vector (Fenno et al., 2014). This system, called INTRSECT, incorporated a modular intron-containing system capable of driving transcription using multiple Boolean logical operations such as AND, NOT, AND NOT, XOR, and NAND (Fenno et al., 2014). A new system following a similar design but using three recombinases (Triplesect), was published while this manuscript was in preparation (Fenno et al., 2020). While extremely elegant, the design for these AND NOT constructs was constrained by the fact that it followed the same internal logic as the AND INTRSECT and Triplesect constructs. As a result of this constraint the currently existing AND NOT system has a couple of shortcomings. The first limitation is that this system adds two introns that make the constructs longer by 194 base-pairs. This is not a serious concern when expressing from a plasmid, but it becomes relevant when dealing with viruses with limited packaging capacity like AAV (Dong et al., 1996). Another concern with the use of introns is that they make the design and cloning of other coding sequences into these vectors more difficult and cumbersome. In this study, we generated an alternative system aimed at solving these constraints. Thus, by design, the ExBoX system has two clear advantages over previous systems: it provides more cargo room and has a simpler design, allowing for replacement of the CDS cassette in a simple cloning step.

The ExBoX system allows for robust expression only in the presence of the respective On switch, which can be reliably inactivated in the presence of the Off switch (Table S1). This is true for both plasmid (Figures 1-4, 7) and viral constructs (Figures 5, 6, 8), as validated by a battery of *in vitro* and *in vivo* experiments in embryonic and postnatal tissues. Very few or no escaper cells still expressed the ExBoX constructs in the presence of the Off recombinase when the On and Off recombinases were active simultaneously (Figures 1-6) or when the Off recombinase was activated by tamoxifen treatment over several days (Figure 7). Considering how efficient the Off switch appeared to be *in vitro* and *in vivo* when using the plasmid vectors (Figures 1-4, 7), the very few escapers observed in the AAV injections (Figures 5, 6, S4) could be explained in part by incomplete or uneven co-infection: the recombinases and the fluorescent markers were driven by independent viral vectors. However, a small but more noticeable number of escapers was observed when FlpO and Cre recombinases were expressed sequentially *in vivo* (Figure 8). It is important to note that in addition to the sequential injection and the aforementioned use of independent viral vectors to drive the recombinases and the fluorescent markers, the titers and ratios for the ExBoX and recombinase vectors were not optimized for these latter experiments. It will be necessary to carefully calibrate these parameters on a case-by-case basis, based on the promoters used to drive expression of the recombinases and the ExBoX constructs, and the stability of the protein of interest in the cells to be studied. Altogether, these experiments demonstrate that the ExBoX system is effective in delivering tight recombinase-dependent expression in a variety of settings.

### A Boolean system to control expression with temporal and cell-type specificity

Boolean logic AND-NOT operations made available by ExBoX can be used to study previously inaccessible subpopulations. In situations where there is no suitable specific driver for a subpopulation of interest, the ExBoX system can be used to drive expression of a gene of interest or reporter in said subpopulation, provided there is a recombinase driver available for a broader neuronal population, and one for the cells to be excluded (Figure 7A, 8). For example, the AND NOT logic of the ExBoX system could be utilized to label or genetically manipulate 5HTR3a^+^;VIP^-^ interneurons in the cortex (Tremblay et al., 2016). This could be done by turning on expression of a CreOn-FlpOff construct with 5*HTR3a-Cre*, while excluding inhibitory neurons expressing *VIP-FlpO* (Figure 7A) (Che et al., 2018; He et al., 2017; Gerfen et al., 2013).

In addition to cell-specificity, these intersectional approaches can be used to achieve temporal control (Figures 7B-F, 8). This is particularly important when studying developmental events or degenerative diseases with defined critical periods. Using ExBoX one could take advantage of the expression of a recombinase to provide cell-type specificity, and a second recombinase to turn Off expression at the desired time (Figures 7B, 8). For example, expression of a gene of interest in the DG can be turned On by injecting ExBoX AAVs at a desired developmental time point into *Grik4-Cre* mice, and later turned Off with Flp, thus creating a window of expression. Our proof of principle experiments demonstrated that ExBoX expression can be shut Off in the order of a few days (Figures 7, 8). Even with a highly efficient system, it is unlikely that protein expression will be turned Off within a few hours, but this will be ultimately dependent on the mRNA and protein stability for each gene of interest. Thus, we think the ExBoX system will be most powerful when studying developmental events that take place over days, but the inclusion of an in-frame PEST sequence might allow for the study of processes with shorter time windows (Rogers et al., 1986).

The tissue- and temporal-specificity conferred by the ExBoX system will make developmental studies that were previously limited by the pre-existing tools possible. With this in mind, we generated ExBoX vectors encoding for neuronal activity and synaptic transmission modulators, which will allow for tight control of these functional manipulations in time and space. These activity-modulating tools, although well characterized in a variety of circuits (see Bando et al., 2016; Burrone et al., 2002; Johns et al., 1999; Lin et al., 2010; Okada and Matsuda, 2008; Priya et al., 2018; Sim and Antolin et al., 2013; Sweeney et al, 1995; Xue et al, 2014 for examples), will need to be validated in each particular setting and/or system. To facilitate studies where the timing of CDS expression is critical, we are currently generating AAV vectors driving expression of a tamoxifen-dependent FlpO (data not shown) (Goodrich et al, 2018). This AAV-FlpOERT2 could be delivered at the same time as the ExBoX AAVs and used to turn Off expression by orally providing tamoxifen at a later time point.

### Expanding the toolkit to study gene function

Due to the ExBoX’s simple design, the system can be easily modified to incorporate alternative and/or additional recombinase systems. For example, inclusion of the Dre-Rox recombinase system, in addition to Cre-lox and Flp-frt, would provide even greater intersectional specificity (Plummer et al, 2015). In addition to Dre-On and Off intersectional AND NOT systems, this could theoretically enable AND-NOT-NOT Boolean logic operations, permitting an even greater expanse of possibilities. In addition, by using other promoters in the AAV constructs this system can be used to drive expression in any cell-type or organ of interest. The broad applicability and exquisite specificity of the system would make the use of these vectors particularly useful for gene therapy applications.

In summary, we designed and generated a new set of viral tools to drive expression using Boolean exclusion logic, ExBoX. ExBoX AAVs allow for tight spatial and temporal control of transcription. These new tools are simple and can be easily modified to express any desired gene of interest. Our design also increases the space available for other coding sequences of interest, even if this increase in capacity is somewhat small (194 bps). As proof of concept, we generated a variety of vectors encoding for neuronal activity and synaptic transmission regulators that will be readily available to the neuroscience community. Based on its simplicity, we believe the ExBoX system will help universalize the use of combinatorial expression approaches, making it a valuable new addition to a series of recently developed intersectional viral systems (Fenno et al., 2014; Fenno et al., 2020; Fan et al., 2020; Sabatini et al., 2021).

Combining these newly developed tools with genetically encoded Cre and FlpO lines, in addition to a growing set of viral vectors, will allow for cell-specific gene expression in a variety of systems.

## Materials and Methods

### Animals

The ICR mouse strain was purchased from Taconic. The *Grik-4-Cre* transgenic mouse strain was purchased from The Jackson Laboratory. Generation of the *Grik4-cre* transgenic mouse line has been described previously (Nakazawa et al., 2002). Mice were housed in a controlled environment maintained at approximately 22°C on a 12-hour light/dark cycle. Mouse chow and water were provided ad libitum. The day of vaginal plug observation was designated as embryonic day 0.5 (E0.5), and the day of birth as postnatal day 0 (P0). For *in vivo* experiments where N=3 at least one female and one male were tested. For N=5, at least 2 males and 2 males were tested. No overt differences in expression of ExBoX constructs were observed between males and females. All animal procedures presented were performed according to the University of California Riverside’s Institutional Animal Care and Use Committee (IACUC) guidelines and approved by UC Riverside IACUC.

### Viral constructs

The CreOn-FlpOff and FlpOn-CreOff ExBoX viral constructs were designed using ApE (https://jorgensen.biology.utah.edu/wayned/ape/) and Vector Builder (vectorbuilder.com) free software. The plasmids were synthesized and assembled by Vector Builder. Plasmid DNA was purified using the Qiagen Maxi Prep kit (Qiagen Cat#10023). AAVs were also packaged by Vector Builder into AAV2/9. pENN.AAV.CamKII 0.4. Cre. SV40 (AAV5) (Addgene viral prep #105558-AAV5; http://n2t.net/addgene:105558; RRID:Addgene_105558), pAAV-CAG-TdTomato (AAV5) (Addgene viral prep#59462-AAV5; http://n2t.net/addgene:59462; RRID:Addgene_59462) and pCAG-FLEX-EGFP-WPRE (AAV9) (Addgene viral prep #51502-AAV9; http://n2t.net/addgene:51502; RRID:Addgene_51502) were obtained from Addgene (Oh, S.W., Harris, J.A., Ng, L., et al., 2014). rAAV5/AAV-EF1α-FlpO-WPRE and rAAV9/CAG-GFP were obtained from the UNC GTC Vector Core. pENN.AAV.hSyn.Cre.hGH was a gift from James M. Wilson (Addgene viral prep # 105555-AAV9; http://n2t.net/addgene:105555; RRID:Addgene_105555). All the ExBoX constructs designed for this study, along with the maps and sequences, were deposited in Addgene.

### *In utero* electroporation

Pregnant ICR mice at E15.5 or E16.5 were anesthetized with isoflurane and an abdominal incision was made to expose the uterus. Pups were visualized through the uterine wall. Plasmids diluted in fast green and phosphate buffered saline (PBS) (3μg/μl) were injected through sharpened glass capillary needles into the lateral ventricle. 5mm paddles were used to deliver five 40V (E15.5) or 37 V (E16.5) pulses of 50 ms each with 950 ms intervals across the hemispheres. Uterine horns were repositioned inside the female and the abdominal cavity was filled with 5x penicillin/streptomycin (pen/strep) in sterile PBS. Pups were collected at E17.5 or postnatally via transcardial perfusion with PBS and 4% paraformaldehyde (PFA) and fixed in 4% PFA for four hours at 4°C, rinsed with PBS, and sectioned coronally at 150 µm on a vibrating microtome (VT100S; Leica). Plasmids were diluted in a 4:1:1 ratio of CreOn-FlpOff-EGFP: pCAG-IRES-Cre: pCAG-IRES-DsRed/pCAG-mCherry-IRES-FlpO or FlpOn-CreOff-TdTomato: pCAG-FlpO: pCAG-EGFP/pCAG-Cre-IRES-GFP. For experiments involving temporal control using tamoxifen-dependent Cre recombinase, the plasmid ratios were 2:1:4:1 of FlpOn-CreOff-TdTomato: pCAG-FlpO: pCAG-ERT2-Cre-ERT2: pCAG-EGFP. For these experiments, pups were birthed and orally gavaged with tamoxifen in corn oil (150 µg/g) or corn oil alone daily for four days starting at P10 until P13 and tissue collected at P23. Immunohistochemistry was performed as described (Polleux & Ghosh, 2002). Nuclei were visualized with DAPI (1 μg/ml) and primary antibodies used were chicken anti-GFP (1:500, Aves Labs Cat# GFP-1010) and rabbit anti-DsRed (1:500, Takara Bio Cat# 632475) with secondary antibodies goat anti-chicken 488 (1:1000, Thermo Fisher Scientific Cat# A-11039) and goat anti-rabbit 546 (1:1000, Abcam Cat# ab60317). Sections were mounted on glass microscope slides with Fluoro-Gel (Electron Microscopy Sciences Cat# 1798502) and imaged with a laser-scanning confocal microscope (Leica SPEII). Fluorescent signals for both GFP and RFP in Figure 4 have been amplified. The fluorescent signals for GFP and RFP in Figure 7E are unamplified and only DAPI was added to visualize nuclei.

### *Ex utero* electroporation

*Ex utero* electroporation was performed similarly as *in utero* electroporation. Pregnant ICR mice at E14.5 were cervically dislocated, uterine horns were removed, and pups were dissected out in Complete HBSS on ice. Plasmids were injected bilaterally and 5 mm paddles were used to deliver five 40 V pulses of 50 ms each with 950 ms intervals across each hemisphere. After electroporation, cortices were removed for primary cortical culture.

### Primary Cortical Neuron Culture

Primary cortical neurons were isolated from E14.5 ICR mouse embryos as described (Kim & Magrané, 2011) using growth media as described (Polleux and Ghosh 2010). 12 mm circular coverslips were treated with 12 M HCL overnight and thoroughly neutralized with deionized water. Acid-treated coverslips were stored in a glass petri dish with 70% to 100% Ethanol. The coverslips were fire-sterilized prior to use and coated with laminin and poly-D-lysine (8.3ug/mL) in sterile water in a 24-well sterile culture plate. *Ex utero* electroporation was performed prior to dissection in Complete Hanks Balanced Salt Solution (HBSS, 2.5 mM HEPES pH7.4, 2.5 mM D-glucose, 1 mM CaCl_2_, 1 mM MgSO_4_, 2.5 mM NaHCO_3_). Neurons were plated at a density of 2.6x10^5^ cells/cm^2^ and maintained in Complete Neurobasal (Neurobasal media, pen/strep, Glutamax, B27) in 5% CO_2_ at 37°C. 50% of the media was exchanged with fresh media 48 hours after plating. Neurons were collected after 72 hours and fixed in 4% PFA for 5 minutes, then rinsed with PBS 3x for 5 minutes. Cells were incubated in blocking buffer (PBS, 1% goat or donkey serum, 0.1% Triton X-100) for 30 minutes at RT, rinsed with PBS 1x, then incubated in primary antibodies (chicken anti-GFP 1:1000, Aves Labs Cat# GFP-1010) and/or rabbit anti-dsRed (1:1000, Takara Bio Cat# 632475) in antibody dilution buffer (PBS, 0.1% goat or donkey serum) at 4°C overnight. The following day cells were rinsed in PBS 4x for 5 minutes and were incubated in secondary antibodies goat anti-chicken 488 (1:1000, Thermo Fisher Scientific Cat# A-11039) and goat anti-rabbit 546 (1:1000, Abcam Cat# ab60317) and DAPI (1 μg/mL) in antibody dilution buffer for 2 hours, then rinsed with PBS 4x for 10 minutes. Coverslips were mounted on glass slides with Fluoro-Gel and imaged with a confocal microscope.

### Neuro2A Cell Culture

Neuro2A cells from ATCC were maintained in DMEM with pen/strep (Gibco Cat#15140-122) and 10% fetal bovine serum and incubated in 5% CO_2_ at 37°C. Cells were visually inspected for contamination daily and routinely tested for Mycoplasma infection using ATCC Universal Mycoplasma Detection Kit (Cat# 30-1012K). Cells were plated on laminin and poly-D-lysine (8.3μg/ml) coated 12 mm coverslips in a 24-well plate at 7.8x10^4^ cells/cm^2^. Cells were transfected 24 hours later with Metafectene PRO (Biontex Cat# T040-1.0) according to the manufacturer’s suggested protocol with a 1:4 DNA to Metafectene ratio. Plasmid ratios were as follows 4:1:1 ratio of CreOn-FlpOff-EGFP: pCAG-IRES-Cre: pCAG-IRES-DsRed/pCAG-mCherry-IRES-FlpO or FlpOn-CreOff-TdTomato: pCAG-FlpO: pCAG-EGFP/pCAG-Cre-IRES-GFP. Cells were fixed 48 hours after transfection with 4% PFA for 5 minutes, rinsed with PBS, stained with DAPI (1 μg/mL) for one hour, then rinsed again 3 times with PBS for 5 minutes. Coverslips were mounted with Fluoro-Gel onto a glass slide and examined with confocal microscopy. GFP and RFP fluorescent signals in Figure 7C,E and Figure S5 are unamplified. In experiments where Neuro2A cells were stained for recombinases (Cre 1:500, EMD Millipore cat# MAB3120; Flp 1:500, Origene cat# TA160030; Figures 1B, 2B, S1, S2) the protocol was as described above, except Triton X-100 concentration was increased to 0.5% for antibody solutions. Secondary antibody used for Cre recombinase was goat anti mouse 647 (1:1000, Life Technologies cat# A21236) and for Flp recombinase goat anti rabbit 647 (1:1000, Invitrogen cat# A21244).

For experiments where tamoxifen was used Neuro2A cells were plated at a density of 2.1x10^4^ cells/cm^2^ 24 hours before transfection and the plasmid ratio was 4:1:1:1 FlpOn-CreOff-TdTomato: pEF1a-Flp: pCAG-ERT2-Cre-ERT2: pCAG-GFP. 4-hydroxytamoxifen in 100% ethanol was added to culture media to 10 μM concentration or equal volume 100% ethanol for vehicle control 48 hours after transfection and replaced daily for three days with fresh 4-hydroxytamoxifen/vehicle, then replaced with regular media as needed until they were collected four days after the last day of tamoxifen.

### Stereotactic Injection

Stereotactic injections were performed as described previously (Osten et al., 2007) in P30 mice. Mice were put under anesthesia for the duration of the procedure using isoflurane. Injections for AAV-FlpOn-CreOff-TdTomato expression experiments were performed in ICR mice targeting the DG of the Hippocampus (coordinates: A/P -1.70 mm; M/L ±1.90 mm; D/V, -1.70 mm). Right hemisphere injections received a 400 nl viral cocktail of AAV-FlpOn-CreOff-TdTomato (6.55x10^12^ GC/mL), rAAV5/EF1α-FlpO-WPRE, pENN.AAV.CamKII 0.4. Cre. SV40, and pCAG-FLEX-EGFP-WPRE in a 1:1:1:1 ratio, whereas the contralateral hemisphere received a 200 nl viral cocktail containing FlpOn-CreOff-TdTomato (6.55x10^12^ GC/mL), EF1α-FlpO-WPRE, in a 1:1 ratio. Titers for commercially available AAVs were as follows: pENN.AAV.CamKII 0.4. Cre. SV40 (2.4 x 10^13^ GC/ml), pCAG-FLEX-EGFP-WPRE (3.3 x 10^13^ GC/ml), pAAV-CAG-TdTomato (1.2 x 10^13^ GC/ml), rAAV5/EF1α -FlpO-WPRE (2.6z x 10^12^ GC/ml), and rAAV9-CAG-GFP (2 x 10^12^ GC/ml). FlpO-negative controls were injected with a viral cocktail of FlpOn-CreOff-TdTomato and pAAV-CAG-GFP. For viral CreOn-FlpOff-EGFP construct experiments, injections were performed in *Grik4-Cre* mice. Right hemisphere injections received 200 nl of desired CreOn-FlpOff construct, whereas the contralateral left hemisphere received a 500 nl viral cocktail containing the construct, rAAV5/EF1α-FlpO-WPRE, and pAAV-CAG-TdTomato in a 2:2:1 ratio (respectively). Viral CreOn-FlpOff-EGFP constructs co-expressing following GOIs were tested: Kir2.1 (3.34x10^13^ GC/ml), Kir2.1-mutant (4.6x10^13^ GC/ml), mNaChBac (2.98x10^12^ GC/ml), mNaChBac-mutant (4.93x10^13^ GC/ml), and TeNT (2.52x10^12^ GC/ml). Three weeks after injections, mice were perfused and fixed with 4% paraformaldehyde O/N at 4°C, rinsed in PBS x1, and sectioned coronally on a vibratome (150 μm). Sections were incubated in 4’,6-diamidino-2-phenylindole (DAPI) and mounted onto slides with Fluoro-Gel fluorescence mounting medium (Electron Microscopy Sciences Cat# 1798502). Expression was observed via confocal microscopy.

### Intracerebroventricular Injections

P1 intracerebroventricular injections were performed as previously described with AAV-CreOn-FlpOff-EGFP or AAV-FlpOn-CreOff-tdTomato in a 1:1 ratio with their corresponding On recombinase AAV (AAV-hSyn-Cre 2.3x1013 GC/ml or rAAV5/EF1α-FlpO-WPRE 2.6 x 10^12^ GC/ml) (Mathiesen et al., 2020; Passini and Wolfe, 2001). In experiments where animals were initially injected at P1 and subsequently injected with Off recombinase into the cortex as adults, 150 nl of viral cocktail was injected at the following coordinates: A/P -1.70 mm; M/L ±1.50 mm; D/V, -0.50 mm. Off recombinases (AAV-hSyn-Cre or rAAV5/EF1α -FlpO-WPRE) were injected 1:1 with pAAV-CAG-TdTomato or rAAV9-CAG-GFP. Controls were injected 1:1 with pAAV-CAG-TdTomato or rAAV9-CAG-GFP and sterile PBS to ensure concentration of viral particles was the same. Tissue was collected 3 weeks after the second injection.

### Image analysis

Neuro2A and primary cortical neuron image analysis was performed with FIJI (Schindelin et al., 2012). GFP+, RFP+, and Cre+, or Flp+ cells were all manually counted. Percentage of transfected cells in which recombinase activity resulted in fluorescent reporter (RFP or GFP) expression were calculated as 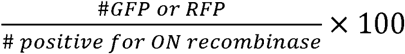 (Figures 1, 2). Images obtained from *in utero* electroporations (Figure 4, 7E,F) were counted manually, avoiding the ventricular zone where it became impossible to distinguish individual cells. Percentage of On cells for primary cortical neuron cultures, Neuro2A transfections for direct fluorescence, and simultaneous AAV injections were calculated as 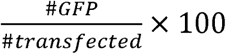 for CreOn-FlpOff and 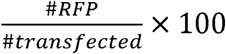 for FlpOn-CreOff (Figures 3, 5, 6, 7C-D, S1A, S2A, S4). The datasets in figures 1, 2, 3, 4, 7, 8, S1, S2, were tested for normality with the Shapiro-Wilk test and QQ plot. A One-way ANOVA with Tukey’s multiple comparisons test was done for Neuro2A transfection, primary neurons, and intrauterine electroporation data (Figures 1, 2, 3D, 4, S1, S2). The data set for some transfections failed to meet normality and thus Fisher’s exact test was performed (Figures 3B, S1, S2). Two-tailed unpaired t-test was performed on data sets in figure 7, 8.

Images from P1 animals intracerebroventricularly injected with AAV-CreOn-FlpOff + AAV-hSYN-Cre that were later cortically injected with rAAV5/EF1a-FlpO-WPRE and pAAV-CAG-TdTomato or pAAV-CAG-TdTomato alone (Figure 8) were quantified by counting the number of On (TdTomato or GFP positive) cells which overlapped with the fluorescent reporter in the second injection of a 100 μm (approximately layer V) that spanned the area of the second injection site and dividing by the number of fluorescent reporter positive cells in the same counting area. For example, CreOn-FlpOff was calculated as 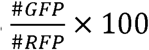. AAV-FlpOn-CreOff images were quantified in the same way 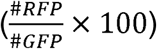. All statistical analyses were performed using GraphPad Prism version 9.2.0 for Windows, GraphPad Software, San Diego, California USA, www.graphpad.com.

## Abbreviations

HA: Hemagglutinin
AAV: Adeno-Associated Virus
DIO: Double-floxed Inverse Orientation
DG: Dentate Gyrus
GOI: Gene Of Interest

## Acknowledgements

We would like to thank Drs. Edward Zagha and Kevin Wright for critically reading the manuscript and providing helpful comments. We would like to thank the University of North Carolina Chapel Hill Gene Therapy Center Vector Core for packaging of viral constructs. pENN.AAV.CamKII 0.4.Cre.SV40 was a gift from James M. Wilson, pCAG-FLEX-EGFP-WPRE was a gift from Hongkui Zeng, and pAAV-CAG-TdTomato and pAAV-CAG-GFP were a gift from Edward Boyden. AAV-EF1α -FlpO-WPRE was a gift from Karl Deisseroth. pCAG-ERT2CreERT2 was a gift from Connie Cepko (Addgene plasmid # 13777; http://n2t.net/addgene:13777; RRID:Addgene_13777). pENN.AAV.hSyn.Cre.hGH was a gift from James M. Wilson (Addgene viral prep # 105555-AAV9; http://n2t.net/addgene:105555; RRID:Addgene_105555)

(Bolded names = Co-first authorship)

