## Supplemental figures for "ExBoX: a simple Boolean exclusion strategy to drive expression in neurons"

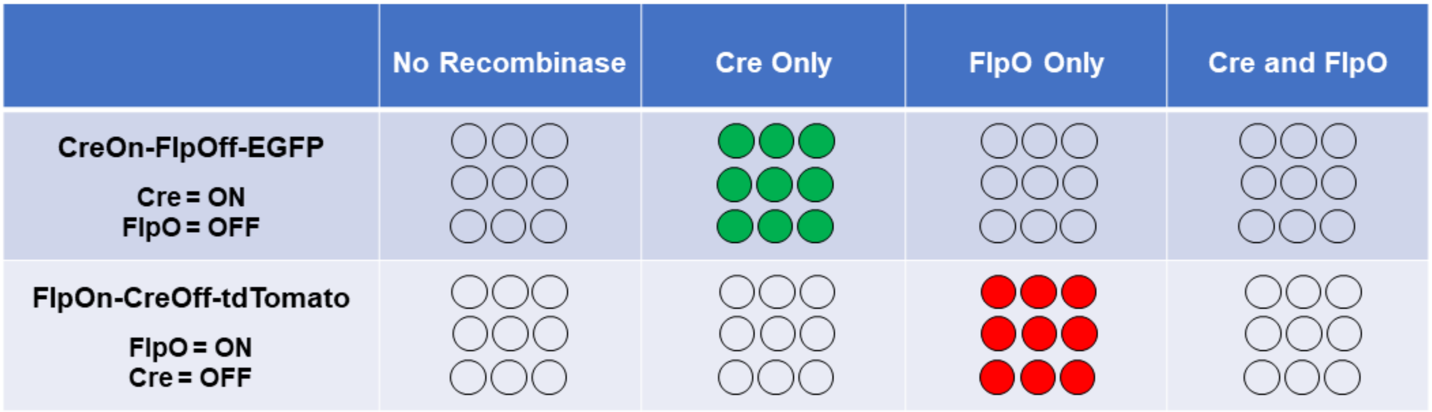


**Table S1. Conceptual basis for the Expression by Boolean Exclusion (AND NOT) system using multiple recombinases.** Expression of a gene of interest (GOI) can be spatially and temporally manipulated using multiple recombinases. In this model, expression of a GOI depends on the presence of an On recombinase and is prevented by the presence of an Off recombinase. Thus, gene expression only occurs in cells where the On recombinase is present AND NOT the Off recombinase. Controlling where and when the On and Off recombinases are expressed provides a high level of specificity for targeting cell populations.

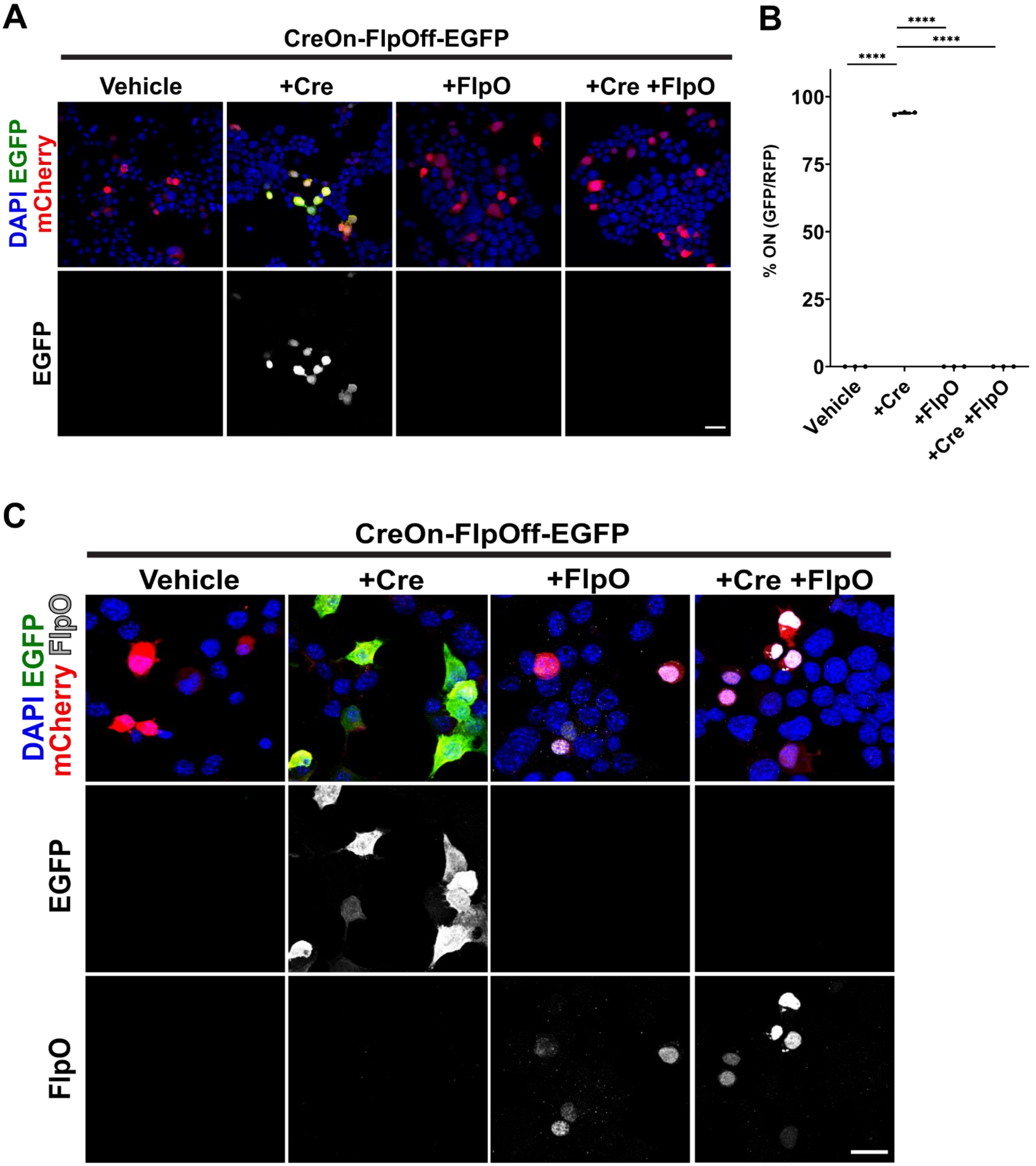

A
CreOn-FlpOff-EGFP
Vehicle
+Cre
+FlpO
+Cre +FlpO
DAPI EGFP mCherry
EGFP
B
****
****
****
% ON (GFP/RFP)
100
75
50
25
0
Vehicle
+Cre
+FlpO
+Cre +FlpO
C
CreOn-FlpOff-EGFP
Vehicle
+Cre
+FlpO
+Cre +FlpO
DAPI EGFP mCherry FlpO
EGFP
FlpO

**Figure S1. Direct fluorescence for CreOn-FlpOff-EGFP in Neuro2A cells and Off recombinase detection. (A)** Unamplified images of Neuro2A cell transfection. **(B)** Quantification of unamplified images shows 93.9%±0.3 of cells transfected with Cre expressing EGFP, while co-transfection with FlpO results in abolishment of EGFP expression. N=3 independent experiments, error bars ±SEM. ANOVA $p<0.001$; Tukey post-hoc test ****$p<0.0001$ vs. all other transfections. **(C)** Immunocytochemical staining for the Off recombinase, Flp, reveals no Flp positive cells remaining in the On condition (EGFP expression). Values given are mean±SEM. Scale bars 30 μm.

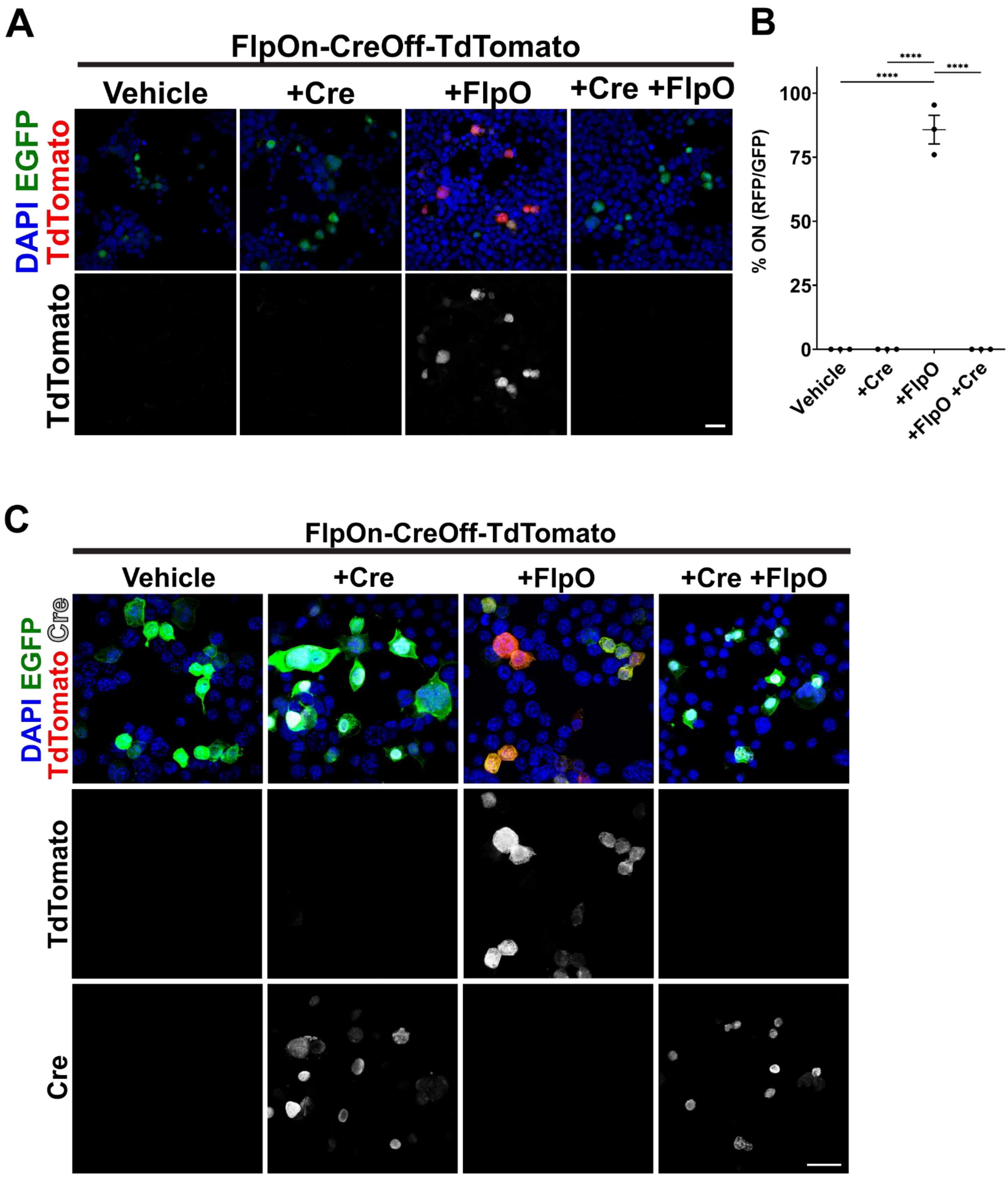
A
FlpOn-CreOff-TdTomato
Vehicle
+Cre
+FlpO
+Cre +FlpO
DAPI EGFP TdTomato
TdTomato
B
****
****
****
% ON (RFP/GFP)
100
75
50
25
0
Vehicle
+Cre
+FlpO
+FlpO +Cre
C
FlpOn-CreOff-TdTomato
Vehicle
+Cre
+FlpO
+Cre +FlpO
DAPI TdTomato EGFP Cre
TdTomato
Cre

**Figure S2. Direct fluorescence for FlpOn-CreOff-TdTomato in Neuro2A cells and Off recombinase immunocytochemistry. (A)** Unamplified images of Neuro2A cell transfection. **(B)** Quantification of unamplified images show expression of reporter in 85.8%±5.6 of transfected cells, while co-transfection with Cre-IRES-EGFP results in abolishment of TdTomato expression. N=3 independent experiments, error bars ±SEM. ANOVA $p<0.01$; Tukey post-hoc test ****$p<0.0001$. Scale bar 30μm. **(C)** Immunocytochemical staining for Cre recombinase reveals no Cre positive cells expressing TdTomato (On condition). Values given are mean±SEM. Scale bar 30 μm.

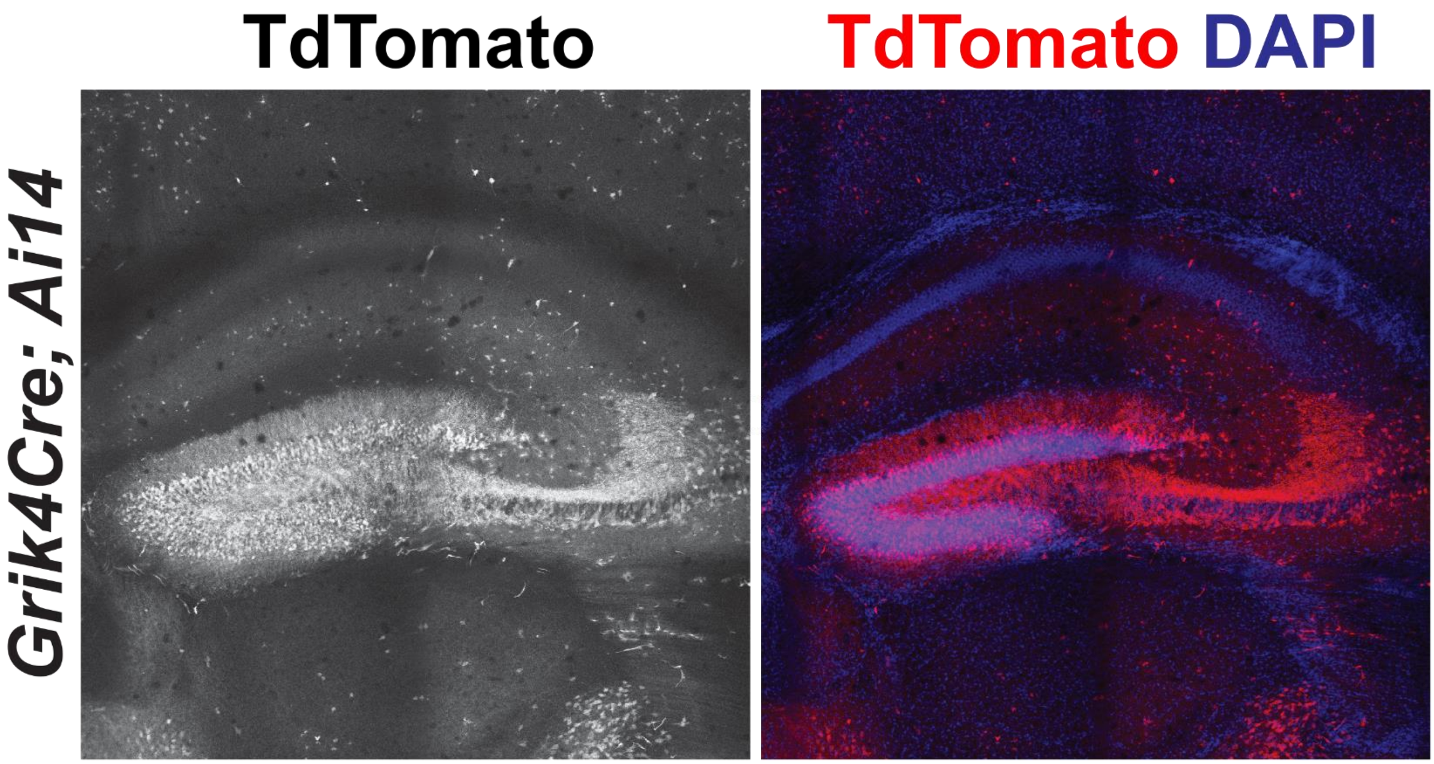


**Figure S3. Expression of Grik4-Cre in the dentate gyrus.** When crossed to a TdTomato reporter mouse (Ai14), robust labeling in the DG is observed.

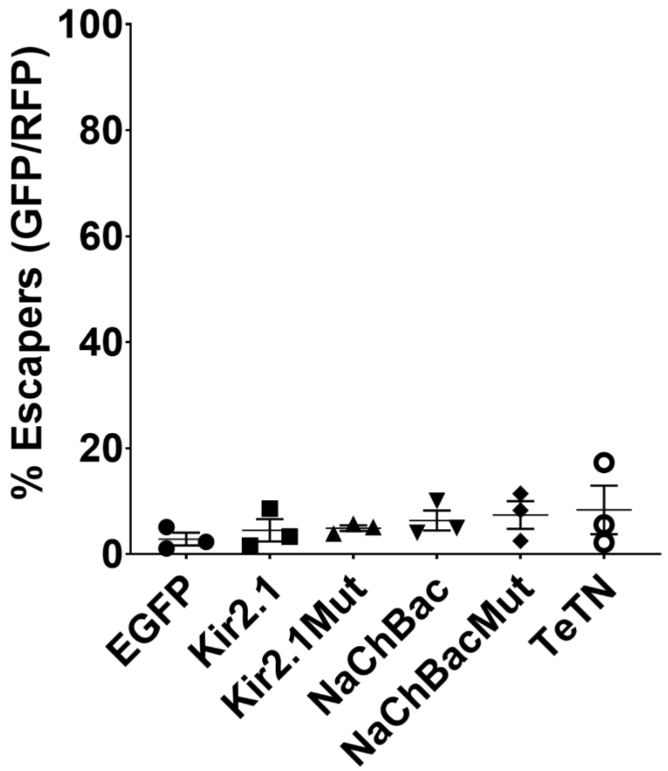


**Figure S4. Quantification of escapers for AAV-CreOn-FlpOff-EGFP constructs expressing activity-modifying GOIs.** AAV-CreOn-FlpOff-EGFP co-injected with AAV-CamkII-Cre (On) and AAV-EF1α-FlpO (Off) resulted in 2.8%±1.2 of cells still expressing EGFP in the presence of the Off recombinase. AAV-CreOn-FlpOff-GOI-EGFP with GOIs Kir2.1 (4.5%±1.2), Kir2.1Mut (4.9%±0.5), mNaChBac (6.3%±1.9), mNaChBacMut (7.4%±2.6), and TeNT (8.3%±4.6) also showed similarly low percentages of escapers when co-injected with the On and Off recombinases (AAV-CamkII-Cre and AAV-EF1α-FlpO, respectively). N=3 independent injections. Values given are mean±SEM.

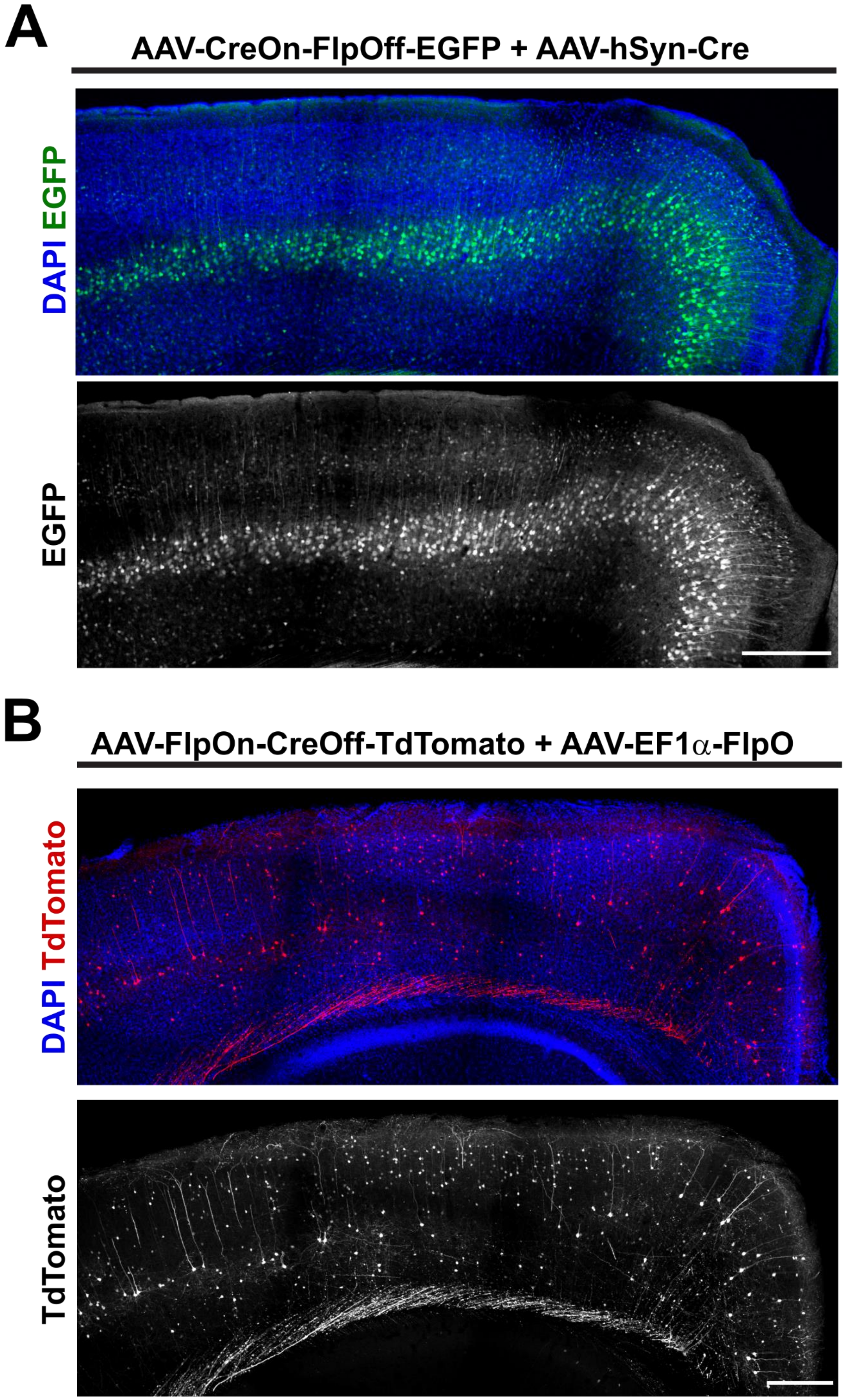
A
AAV-CreOn-FlpOff-EGFP + AAV-hSyn-Cre
DAPI EGFP
EGFP
B
AAV-FlpOn-CreOff-TdTomato + AAV-EF1α-FlpO
DAPI TdTomato
TdTomato

**Figure S5. Unamplified images of ExBoX AAVs.** Unamplified images of AAV-CreOn-FlpOff-EGFP **(A)** and AAV-FlpOn-CreOff-TdTomato **(B)**. Animals were injected intracerebroventricularly at P1 and tissue was collected for visualization 3 weeks later. Imaging was done after perfusion and 4% PFA postfix for 4 hours.
